# Shared Representation Discovery for Multi-Subject Neural Data Analysis

**DOI:** 10.64898/2026.08.18.745594

**Authors:** Felix H. Taschbach, Marcus K. Benna

## Abstract

Neural recordings from different individuals vary substantially even when behavior is broadly shared. The sampled neurons differ, and the same behavior occurs at different times. Standard cross-subject analyses rely on matched time points or anatomical correspondence, which excludes many datasets. We recently introduced Shared Representation Discovery (ShaReD), which identifies neural-behavioral relationships conserved across subjects by learning a shared behavioral projection together with subject-specific neural projections. Here we develop and benchmark this method using synthetic, primate, and rat data. On synthetic data, ShaReD recovers common structure across noise levels, sample sizes, and subject counts, and separates components confined to different groups of subjects. In non-human primate motor cortex, ShaReD identifies kinematic representations that generalize across individuals and across reaching tasks with different movement statistics. In rats navigating a spatial alternation task, ShaReD isolates behavior-aligned directions within the CA1-to-PFC communication subspace. ShaReD thus extends multi-subject analysis to datasets in which comparable behaviors occur without cross-subject temporal correspondence.

## Introduction

Behavior provides a direct and readily available set of observables influenced by an animal’s brain activity, and the relationship between neural activity and measured behavioral variables is a primary quantity of interest in neuroscience. When sufficient amounts of data are available we can study this relationship in individual subjects, but pooling across animals allows us to determine whether a neural code generalizes rather than reflects one individual’s idiosyncrasies, and it adds statistical power when any one recording is short or has few neurons. Analyses of this kind usually assume that every animal encodes behavior in the same way, but this may not be the case in general.

Even when that assumption is approximately correct, the observed activity patterns of neural populations vary across individuals. The sampled neurons are not the same, and the trajectories through firing rate space differ. In contrast, even though any given behavior can occur at different times, the behavioral variables observed are often the same across individuals. For example, position, velocity, posture, task state, choice, and reward are measured in the same coordinates across subjects even when their neural implementation differs.

Comparing such populations requires a correspondence between subjects, which existing methods define in various ways. Hyperalignment [1, 2] and Shared Response Models [3] learn transformations into a common neural space using time-locked stimuli. Canonical correlation analysis aligns motor cortical dynamics across animals by matching reaching conditions [4]. Contrastive approaches such as CEBRA [5] learn joint neural-behavioral embeddings from temporal labels. Dynamic time warping relaxes exact time locking but assumes that behavioral elements occur in the same order [6]. These assumptions are appropriate for controlled, trial-based experiments with matched timing. They do not extend to more naturalistic datasets in which subjects produce related behaviors at different times, for different durations, and in a different order [7].

The question of interest is often whether different neural populations carry the same behavioral signal. When the recorded neurons differ and no time-locked stimulus anchors them, this question cannot be answered by aligning the neural activity directly, because the alignment is underdetermined. The correspondence thus has to come from the behavior, which is measured in a common coordinate system across subjects. The neural-behavioral relationships, rather than the neural trajectories, can then be compared across individuals to address neural encoding and decoding questions.

## Background

Shared Representation Discovery (ShaReD) is designed to facilitate exactly this comparison and to extract common neural-behavioral relationships present in multiple subjects. For *K* subjects, let **X***_k_ ∈* R*^Tk ×Nk^* denote the activity of *N_k_* recorded neurons in *T_k_* time bins, and **Y***_k_ ∈* R*^Tk ×B^* the *B* behavioral variables recorded at those same time points. ShaReD learns a subject-specific neural loading (or projection) **a***_k_ ∈* R*^Nk^* and a shared behavioral loading **b** *∈* R*^B^* (Fig. 1A). Each neural population is then projected to **X***_k_***a***_k_* and each behavioral recording to **Y***_k_***b**, which corresponds to one component describing a particular relationship between correlated neural and behavioral quantities. Further components, each describing a different shared relationship, are extracted sequentially (see Methods). The number of neurons and time points can vary across subjects, while the behavioral variables are measured in a common coordinate system [8]. This procedure thus places its central constraint on the behavioral side, where the experimental recordings provide structure common to all subjects. Because neural representations may differ across animals, each subject has its own neural loading. The behavioral variables, by contrast, are shared by design, so all subjects share one behavioral loading. When subjects encode different combinations of behavioral features, the shared **b** describes the direction in behavioral feature space most consistent across subjects rather than any single subject’s encoding. ShaReD therefore searches for combinations of behavioral variables whose neural correlates recur across subjects.

**Figure 1:**
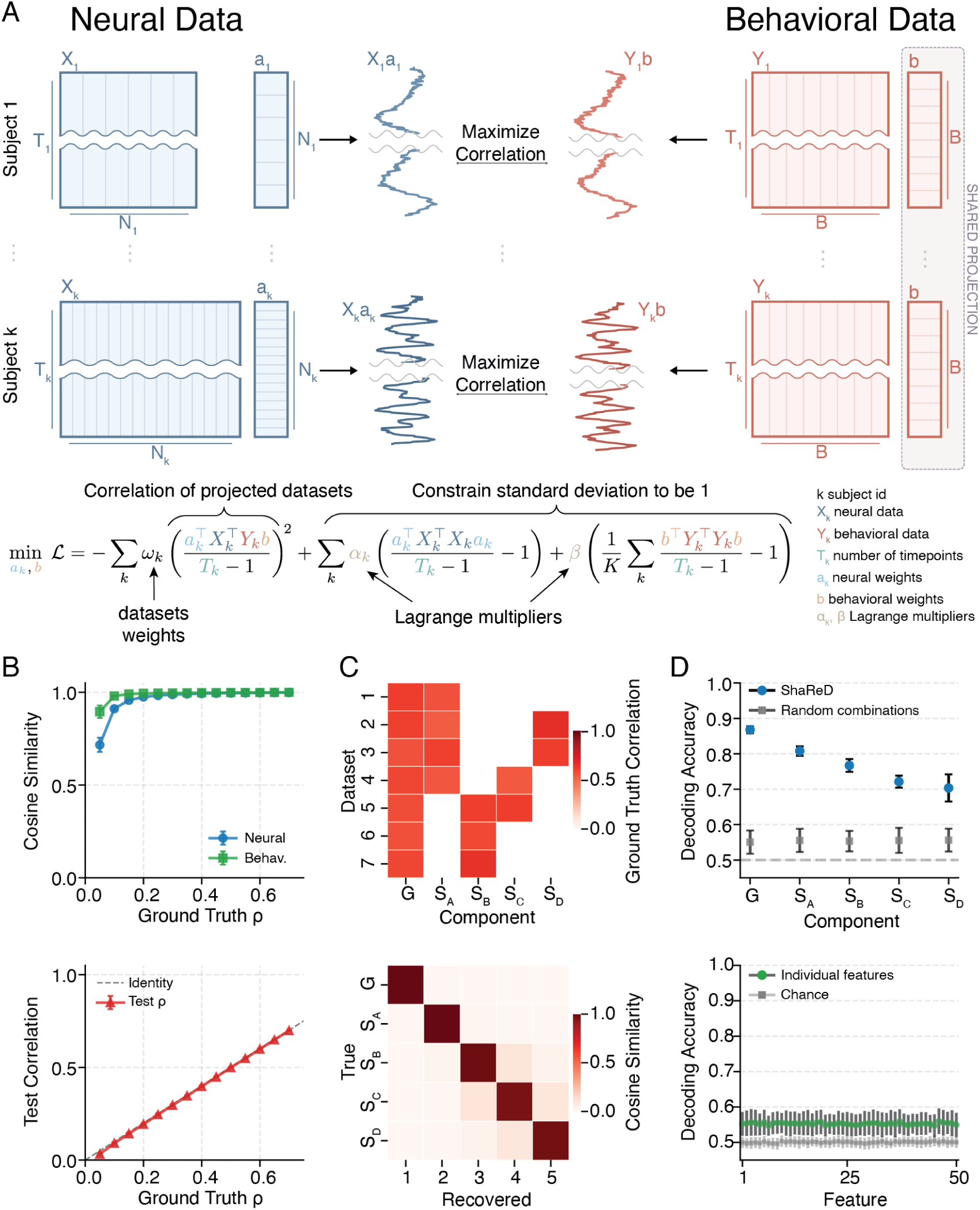
ShaReD method and synthetic data validation. **(A)** Schematic of Shared Representation Discovery. For *K* subjects with paired neural (**X**, blue) and behavioral (**Y**, red) data, ShaReD identifies subject-specific neural projections (**a***_k_*) and a single shared behavioral projection (**b**) that maximize the sum of squared correlations between projected time series. Neural data matrices can have different numbers of neurons (*N_k_*) and time points (*T_k_*), while the set of measured behavioral features (*B*) is consistent across subjects. The objective function (bottom) weights each subject’s contribution (using coefficients *ω_k_*), while Lagrange multipliers (*α_k_*, *β*) enforce unit-variance constraints. **(B)** Component recovery on synthetic data (6 subjects with 14 simultaneously present components with ground truth correlations from 0.05 to 0.70 spaced in equal increments of 0.05). *Top*, cosine similarity between true and recovered projections remains high (*>* 0.9) for components with ground-truth correlation *ρ ≥* 0.1. Behavioral projections (green) are recovered more accurately than neural projections (blue). *Bottom*, cross-validated test correlations closely track the identity line, evidence of generalization without overfitting. Error bars indicate SD across 100 repetitions. **(C)** Recovery of components with different sharing structure across subjects. Each component is defined by its support, the set of subjects that encode it, rather than by its strength. All five components are matched in aggregate correlation strength (*ρ̄* = 0.60), so they differ only in which subjects share them. *Top*, ground-truth encoding structure showing 1 global component (G), 2 subgroup components (S*_A_*, S*_B_*), and 2 partially overlapping components (S*_C_*, S*_D_*) with distinct sharing patterns across 7 subjects. *Bottom*, cosine similarity between true and recovered behavioral projections shows clean block-diagonal structure. ShaReD identifies components at every level of sharing without prior specification. **(D)** Pseudopopulation decoding. *Top*, decoding accuracy for binarized ShaReD component states from pseudopopulations constructed by sampling neurons across subjects (blue circles). The global component (G) achieves the highest accuracy (*∼*0.87), with subgroup components ranging from *∼*0.70 to *∼*0.81. Decoding from random linear combinations of behavioral features (gray squares) is substantially lower (*∼*0.55) across all components. The ShaReD axes capture neurally accessible structure beyond what arbitrary behavioral mixtures provide. Dashed line indicates chance (0.5). *Bottom*, decoding individual behavioral features from pseudopopulations yields only marginally above-chance accuracy (*∼*0.55–0.56, green) compared to chance (*∼*0.50, gray). The cross-subject decodability in the top panel therefore reflects shared component structure rather than information in any single feature. Multiple components are extracted sequentially via deflation (see Methods). Error bars indicate SD across 100 repetitions.

This approach leads to a multi-subject analogue of canonical correlation analysis (CCA) [9]. Evidence is pooled through one behavioral loading rather than fit independently per subject, so even a subject with a short recording or few neurons can be usefully combined with data from the rest of the subjects. The objective function is

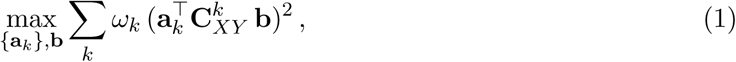

where 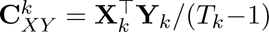 is the neural-behavioral cross-covariance for subject *k*, and *ω_k_* weights each subject’s contribution. Each projected time series is constrained to have unit variance, with the behavioral constraint pooled across subjects (see Methods).

The squared objective addresses a multi-subject failure mode of linear correlation objectives. When subjects share two components with different relative strengths, a linear objective can prefer a mixed direction over either of the unmixed components. Squaring the correlations makes the objective an affine function of the mixing between the two components, and the optimum lies at an unmixed component unless the aggregate evidence for the two components is exactly tied. A formal comparison of the linear, squared, and higher-power objectives is given in Supplementary Note 1.

ShaReD was previously used to extract a single shared kinematic axis in a study of motor learning [8]. Here, we provide the full methodological treatment of ShaReD. We validate the method on synthetic data with known ground truth, apply it to macaque motor cortical recordings during reaching, and extend it to simultaneous hippocampal-prefrontal recordings in rats. The macaque analysis tests whether shared kinematic components generalize across animals and across reaching tasks, and it benchmarks behavior-based alignment against neural alignment with and without temporal correspondence. The rat analysis uses ShaReD inside a CA1-to-PFC communication subspace to separate behavior-aligned structure from residual signals, including prospective choice encoding.

## Results

### ShaReD recovers shared structure in synthetic data

To validate ShaReD against known ground truth, we generated datasets from a linear latent factor model with the projections and per-component correlations set directly (Methods, Supplementary Note 3). The test is whether ShaReD recovers the ground-truth projections and whether the recovered correlations generalize rather than overfit. Each dataset had 6 subjects with 50 neural features, 50 behavioral features, and 20,000 training samples per subject, and contained 14 components present simultaneously at target correlations from 0.05 to 0.70. At this sample size, ShaReD recovered behavioral and neural projections with cosine similarity above 0.9 for components with true correlation *ρ ≥* 0.1 (Fig. 1B, top). Behavioral projections were recovered more accurately than neural projections, reflecting the many-to-one mapping from neural implementations to behavioral outputs. Cross-validated correlations on held-out data tracked the target correlations across the full range of signal strengths (Fig. 1B, bottom).

Recovery scaled with sample size, signal strength, and subject count, and degraded with noise (Supplementary Fig. S3). Components above *ρ* = 0.5 were recovered consistently from 500 samples, components between 0.2 and 0.5 required approximately 2,000 samples, and the weakest component (*ρ* = 0.05) was not reliably recovered at any tested sample size.

ShaReD did not require knowledge of which subjects shared which components. We generated 7 subjects with 5 components matched in aggregate correlation strength (*ρ̄* = 0.60) but differing in their support, comprising 1 global component shared by all subjects, 2 subgroup components each confined to a contiguous block, and 2 pair components each shared by 2 subjects (Fig. 1C, top). The recovered behavioral projections separated all 5 components and reproduced the expected block structure, recovering the sharing structure from the data alone (Fig. 1C, bottom).

The recovered behavioral axes provided a reference frame for pooling neural data across subjects. We projected behavioral data onto each recovered component direction, binarized the resulting values by median split, and decoded the labels from pseudopopulation neural activity using logistic regression. The global component was decoded most accurately (*∼*0.87), with subgroup components ranging from *∼*0.70 to *∼*0.81 (Fig. 1D, top). Decoding from random combinations of behavioral features in the same pseudopopulations was substantially lower (*∼*0.55), and decoding individual behavioral features yielded only marginally above-chance accuracy (*∼*0.55–0.56, Fig. 1D, bottom).

Because the data were generated from ShaReD’s own linear model, this analysis is a recovery test rather than an independent validation.

### ShaReD identifies consistent motor representations across animals

We asked whether ShaReD recovers kinematic components that generalize across subjects. We analyzed data from 3 macaques performing either a center-out (CO) task with 8 fixed radial targets or a random target (RT) task with unpredictable target positions (Fig. 2A). Because neuron identities differed across sessions, including sessions from the same animal, each session was treated as a separate dataset. The CO analysis used 73 sessions and the RT analysis used 26 sessions, with 33 to 295 neurons per session. The RT task had greater behavioral variability than the CO task (mean position variance 17.2 *±* 4.4 cm^2^ for CO and 32.6 *±* 2.0 cm^2^ for RT, *P <* 10*^−^*^13^, Mann–Whitney *U*).

**Figure 2:**
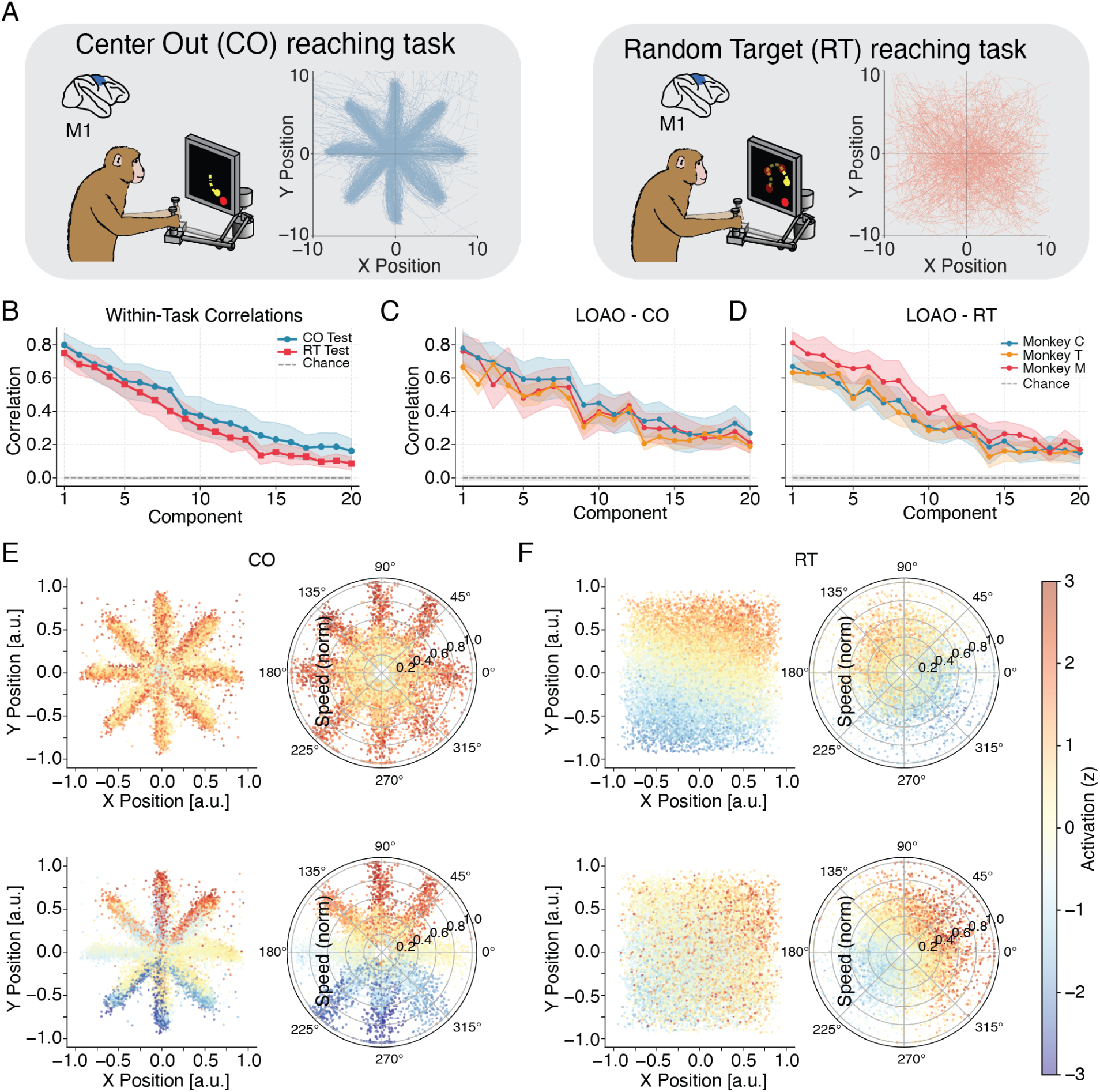
ShaReD identifies common kinematic components in macaque motor cortex. **(A)** Task design. Monkeys performed either a center-out (CO) task with 8 fixed radial targets and stereotyped trajectories, or a random target (RT) task with unpredictable positions and variable movement patterns. Neural activity was recorded from primary motor cortex (M1) during both tasks. Task schematics adapted from [4]. **(B)** Within-task cross-validated neural-behavioral Pearson correlations for CO (blue, *n* = 73 sessions) and RT (red, *n* = 26 sessions) tasks across 20 extracted components. Shaded regions indicate SD across sessions. Both tasks yield multiple components substantially above chance (gray dashed line). CO correlations exceed RT correlations.**(C)** Leave-one-animal-out (LOAO) validation for the CO task. Behavioral projections learned from 2 animals generalize to held-out sessions from the third animal. Colors indicate the held-out monkey (C in blue, T in orange, M in red), and shaded regions indicate SD across held-out sessions. All animals show cross-validated neural-behavioral Pearson correlations well above chance. ShaReD identifies consistent rather than animal-specific encoding. **(D)** Leaveone-animal-out validation for the RT task, as in (C). **(E, F)** Visualization of the first two ShaReD components in behavioral space for the CO task (E) and RT task (F). Each row corresponds to one component. Left subpanels show hand position (*x* versus *y*) colored by projected component value (*z*-scored activation, blue low, red high). Right subpanels show polar movement-direction tuning, where angle indicates movement direction and radius indicates normalized speed. In CO, the first component is a radial signal tuned to speed and distance from center, and the second component is sensitive to vertical position. In RT, the first two components are approximately tuned to vertical and horizontal position. The same kinematic features recur across tasks, though not necessarily in the same component.

Behavioral features included hand position (*x*, *y*), speed, and distance from workspace center at 31 time lags spanning *±*500 ms, giving 124 features per time point. Feature selection details are in the Methods, with representative ablations shown in Supplementary Fig. S10. All 3 kinematic variables contributed, with the dominant variable differing by task. Adding acceleration or jerk did not improve performance, and reducing the temporal window from *±*500 ms to *±*333 ms reduced performance by approximately 0–1% (with a larger *∼*1–3% drop at *±*167 ms).

ShaReD extracted multiple neural-behavioral components in both tasks (Fig. 2B). First component correlations were similar across tasks (0.80*±*0.07 for CO and 0.75*±*0.07 for RT). Across the first 20 components, the mean correlation was higher for CO than for RT (0.41 *±* 0.08 versus 0.34 *±* 0.06, mean *±* SD across components). The difference held per session, with each session summarized by its mean correlation over the 20 components (Mann–Whitney *U* = 1412, *P* = 2.4 *×* 10*^−^*^4^, *n* = 73 CO and 26 RT sessions), reflecting the lower behavioral variability of the CO task. We restrict the analysis to the leading 20 components throughout. A per-component significance test finds more than 20 components individually significant (Supplementary Fig. S13), so 20 is a conservative cap rather than the limit of recoverable structure.

Behavioral projections learned from 2 animals remained predictive in the third. ShaReD was fit to sessions from two animals, and the learned behavioral projections were evaluated on the held-out animal. For each held-out session, neural projections were fit using 10-fold cross-validation while the behavioral projections were fixed. First component correlations ranged from 0.63 to 0.81 across held-out animals and tasks (Fig. 2C,D). The learned behavioral directions were not animal-specific.

The leading components captured interpretable kinematic features, though the features carried by a given component index differed between tasks (Fig. 2E,F and Supplementary Figs. S4, S5). In CO, the first component was a radial signal tuned to speed and distance from center with no directional preference, and the second was tuned to vertical hand position. In RT, the first two components were approximately tuned to vertical and horizontal position. They showed directional tuning in the polar plots, modulated by speed. Temporal weight profiles of the leading components varied smoothly across the delay window, while weaker, higher-index components were less specific to a single feature. Because a feature was not tied to a fixed component index across tasks, we matched components by their behavioral weights rather than by their order when comparing tasks, and how well each component transferred tracked this weight similarity (Fig. 3B).

**Figure 3:**
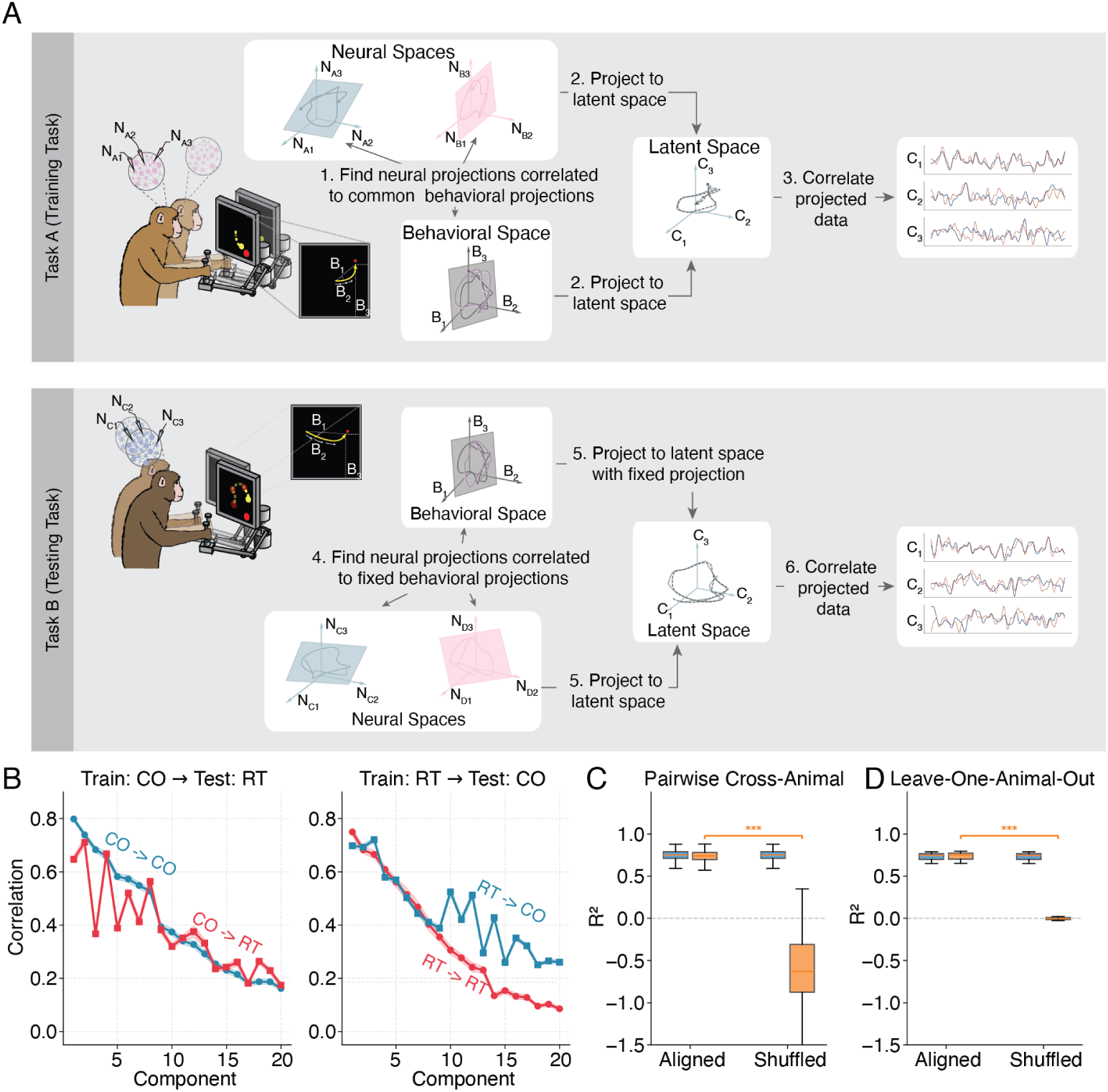
ShaReD components transfer across tasks and tolerate disrupted temporal correspondence. **(A)** Schematic of cross-task validation procedure. *Training phase (Task A)*, ShaReD is applied to multiple animals performing one task to identify shared behavioral projections and subject-specific neural projections. *Testing phase (Task B)*, using the fixed behavioral projections from training, new neural projections are learned for each animal in the test task, and cross-validated correlations are computed to assess generalization. Task schematics adapted from [4]. **(B)** Cross-task generalization results showing Pearson correlation across sessions. *Left*, components trained on CO task (CO*→*CO, blue) compared with their transfer to RT task (CO*→*RT, red). *Right*, components trained on RT task (RT*→*RT, red) compared with their transfer to CO task (RT*→*CO, blue). Components transfer in both directions. Transferred correlations track the baseline difficulty of the testing task rather than the training task. CO-tested correlations tend to exceed RT-tested correlations regardless of training source. Shaded bands indicate SEM across sessions. **(C, D)** Effect of temporal misalignment on cross-animal position decoding (*R*^2^). ShaReD (blue) and GCCA-based alignment (orange) achieve similar performance with temporally aligned data. When temporal correspondence is disrupted by permuting one session’s target labels, GCCA performance collapses while ShaReD is unaffected, since its projections depend only on within-session neural-behavioral correspondence. **(C)** Pairwise cross-animal decoding (*n* = 1,300 session pairs). Boxes show the interquartile range with the median, and whiskers extend to 1.5 times the IQR. **(D)** Leave-one-animal-out generalization (*n* = 73 sessions, 3 held-out animals). Boxes show the interquartile range with the median, and whiskers extend to 1.5 times the IQR. The GCCA aligned-versus-shuffled change in decoding *R*^2^ was assessed by a clustered sign-flip permutation test (10,000 permutations). The null flips the sign of each session cluster’s aligned-minus-shuffled difference. In (C) each cross-animal pair carries two session memberships and is flipped by the product of its two sessions’ signs, and in (D) each observation is a single session (Methods). *^∗^P <* 0.05, *^∗∗^P <* 0.01, *^∗∗∗^P <* 0.001.

The shared behavioral constraint preserved most of the single-session neural-behavioral structure. Within-session ShaReD correlations were comparable to unconstrained single-session CCA (Supplementary Fig. S7). Behavioral projections learned independently by single-session CCA were unstable across sessions, even after aligning components across sessions by optimal (Hungarian) matching [10] and sign correction, with mean absolute cosine similarity across 20 components of 0.50 (CO) and 0.49 (RT) (Supplementary Fig. S8). ShaReD’s shared projections were instead reproducible across cross-validation folds, with mean absolute cosine similarity above 0.99 for the first *∼*12 components (Supplementary Fig. S11). Deflating ShaReD components reduced subsequent CCA correlations. ShaReD therefore captured the dominant neural-behavioral covariance also found by unconstrained CCA while imposing a common behavioral coordinate system (Supplementary Fig. S9). Results were stable across the number of delays, optimization iterations, the smoothness penalty, and the objective power (quadratic vs. quartic) (Supplementary Fig. S12).

### ShaReD components generalize across reaching tasks

Behavioral projections learned in one task remained predictive in the other. ShaReD was fit to all sessions from one task to identify shared behavioral projections, which were then fixed. For each session in the other task, neural projections were fit using the training folds, and correlations were evaluated on held-out data (Fig. 3A).

Transfer occurred in both directions (Fig. 3B). Conditions tested on CO reached similar correlations whether trained within task or transferred (CO*→*CO 0.41 *±* 0.08 and RT*→*CO 0.44 *±* 0.07, mean *±* SD across 20 components), as did conditions tested on RT (RT*→*RT 0.34 *±* 0.06 and CO*→*RT 0.38 *±* 0.06). Both CO-tested conditions exceeded both RT-tested conditions regardless of the training task, so transfer was governed by the difficulty of the test task rather than the training task. The behavioral projections learned in one task therefore stayed predictive in the other, evidence that the shared components generalize across the two tasks.

Transfer was component-specific. Behavioral weight similarity after Hungarian matching [10] ranged from 0.25 to 0.82 across the first 10 components (Supplementary Fig. S6). Transfer from CO to RT (the task with higher variability) was weaker for components whose behavioral weights were less similar between tasks, most clearly for component 3, the least similar pairing (Figs. 3B, S6A).

### Behavior-based alignment is robust to disrupted temporal correspondence

Methods that align subjects through neural activity require a sample-to-sample correspondence across subjects. That correspondence generally follows from an auxiliary variable, usually time within a trial under the assumption that all subjects express the same behavior at the same trial-relative moment, which allows the matched time points to be aligned. ShaReD aligns through each subject’s own neural-behavioral relationship instead, so no such correspondence is needed. We compared the two approaches in regimes where this time-based correspondence either held or was disrupted, using GCCA-based neural alignment as a comparison baseline [4, 11] (Fig. 3C,D).

With temporally aligned reaching movements, the two approaches achieved similar cross-animal position decoding (ShaReD *R*^2^ = 0.75 *±* 0.07, GCCA *R*^2^ = 0.74 *±* 0.07, mean *±* SD, *n* = 1,300 cross-animal session pairs, Fig. 3C). We then broke the correspondence between paired sessions by permuting the target labels of one session in each pair. GCCA decoding dropped sharply when shuffled, to *R*^2^ = *−*0.63 *±* 0.37 (aligned-minus-shuffled difference *P <* 10*^−^*^4^, clustered sign-flip permutation test, 10,000 permutations), while ShaReD was unchanged (*R*^2^ = 0.75 *±* 0.07).

Leave-one-animal-out tests gave the same result (Fig. 3D). With aligned samples, ShaReD and GCCA performed similarly (ShaReD *R*^2^ = 0.729 *±* 0.007, GCCA *R*^2^ = 0.734 *±* 0.007, mean *±* SD over 6 cross-validation folds, *d_z_* = *−*0.60). After the correspondence was disrupted, GCCA performance fell to chance (*R*^2^ = *−*0.021 *±* 0.007, aligned-minus-shuffled difference *P <* 10*^−^*^4^, clustered sign-flip permutation test with 10,000 permutations), while that of ShaReD remained unchanged (*R*^2^ = 0.729 *±* 0.007). ShaReD therefore matches neural-alignment methods when temporal correspondence between data points across different sessions is available, and is unaffected by its absence.

### ShaReD isolates behavior-aligned structure within the hippocampal-prefrontal communication subspace

Reduced-rank regression (RRR) identifies low-dimensional activity patterns in one brain region that predict activity in another [14]. CA1 and PFC both encode spatial variables during navigation, so a CA1-to-PFC prediction can reflect direct signaling between the areas, behavioral variables encoded in both, or a mixture of the two, and no correlation-based method separates these. We can, however, remove the part of the prediction tied to specific measured behaviors. We used ShaReD to find the directions within the CA1-to-PFC communication subspace whose activity is predicted by the same behavioral features in every rat, and asked how much of the prediction they account for and what remains once they are removed.

We first fit RRR independently in each of 7 rats navigating a W-track in an alternation task and fixed the CA1-to-PFC communication subspace at 11 dimensions for all animals, the median of the per-animal ranks selected by a one-standard-error (1-SE) rule (Supplementary Fig. S14). We then applied ShaReD across animals to these communication-subspace latents (Fig. 4A,B). Six of the 11 ShaReD components had significant cross-validated neural-behavioral correlations (refit-shuffle permutation test with Benjamini–Hochberg FDR correction, *q <* 0.05, with components 1 to 5 reaching *p*_FDR_ = 0.004 and the sixth *p*_FDR_ = 0.040, Fig. 4C, Supplementary Fig. S15).

**Figure 4:**
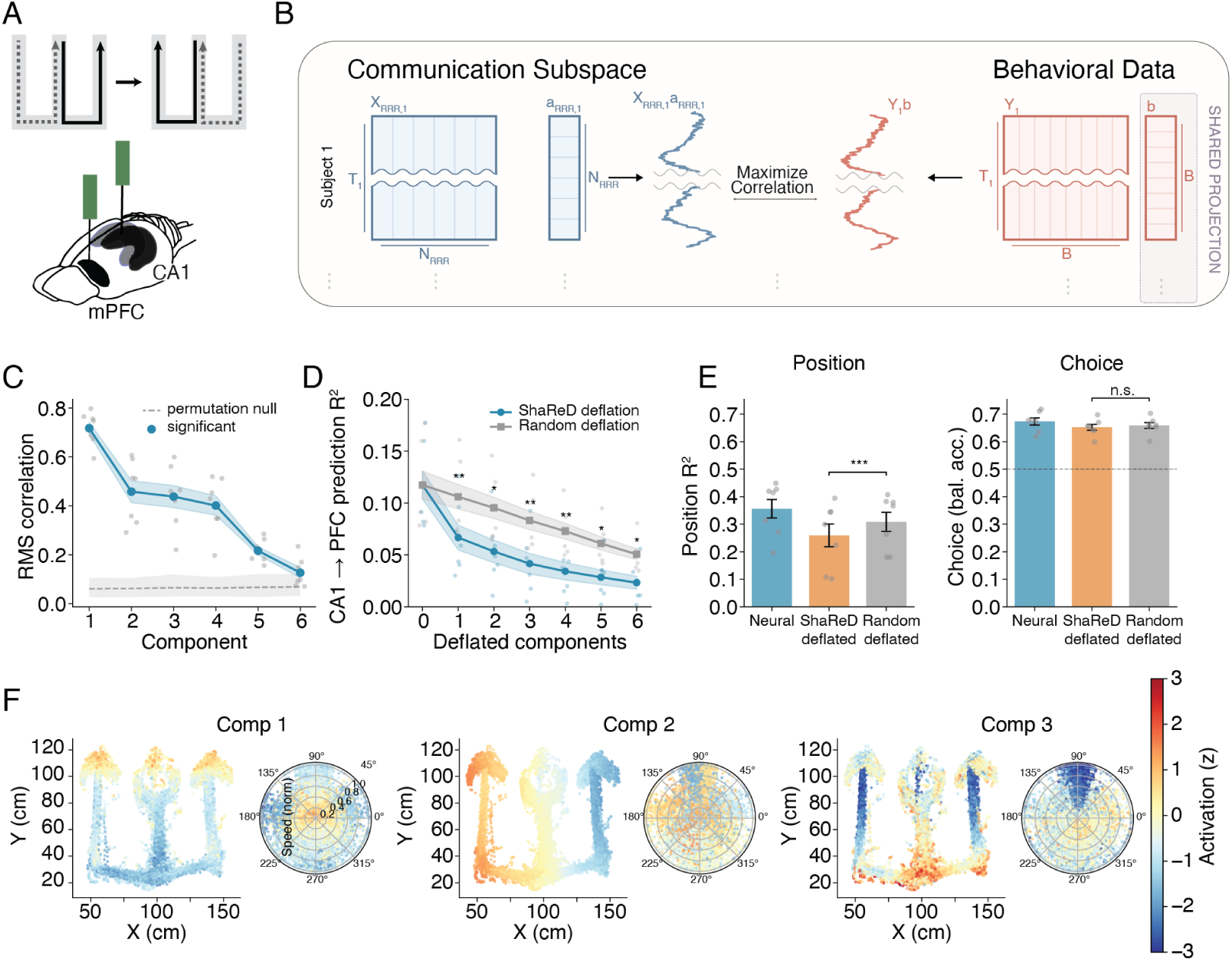
Behavior-aligned structure within the hippocampal-prefrontal communication subspace. **(A)** Experimental setup adapted from [12, 13]. Rats navigated a W-shaped track while simultaneous recordings were obtained from hippocampal CA1 and medial prefrontal cortex (PFC). **(B)** Two-stage analysis schematic. Reduced-rank regression (RRR) independently identifies each animal’s CA1-to-PFC communication subspace (left). ShaReD then finds a shared behavioral projection across animals within that subspace (right) that maximizes the correlation between projected neural and behavioral time series. **(C)** Cross-validated RMS correlation between neural and behavioral projections for the 6 significant ShaReD components extracted from the communication subspace, shown against the permutation null (gray dashed line). The blue line shows mean ± SEM and gray dots show individual animals. **(D)** CA1-to-PFC prediction *R*^2^ after progressively deflating ShaReD-identified behavioral dimensions (blue) versus random dimensions of matched rank (gray). ShaReD deflation reduces prediction significantly more than random deflation. Stars indicate significant differences (Bonferroni-corrected). Shaded bands indicate SEM across 7 rats. **(E)** Dissociation between position and choice information. In both panels the 6 significant ShaReD components are deflated, with random deflation of matched rank (6 dimensions) as a control. *Left*, position decoding (*R*^2^) from the full CA1 population (blue), after ShaReD deflation (orange), and after random deflation (gray), on both phases of outbound trials. ShaReD deflation reduces position *R*^2^ significantly more than random deflation (*Z* = *−*5.73, *P* = 1 *×* 10*^−^*^8^, signed Liptak–Stouffer, Bonferroni-corrected for 2 comparisons). *Right*, choice decoding (balanced accuracy) on stem-phase data of outbound trials with causal-only delays. ShaReD deflation does not significantly reduce choice decoding relative to either the full population or random-deflated controls, scored by balanced accuracy with balanced class weights so the metric is robust to label imbalance (Methods). Deflating the common behavioral components thus reduces position information substantially more than random deflation and has no comparable effect on prospective choice. Dashed line indicates chance (0.5). Significance brackets show per-animal empirical-null tests combined by signed Liptak–Stouffer with Bonferroni correction for 2 comparisons. Error bars indicate SEM across 7 rats. *^∗∗∗^P <* 0.001*,^∗∗^ P <* 0.01*,^∗^ P <* 0.05, n.s. not significant. **(F)** Spatial and directional tuning of the first 3 ShaReD components for an example animal (ER1). Scatter plots show component activation as a function of position on the W-track (color scale, *z*-scored). Polar plots show head-direction tuning. Components capture distinct spatial features. *n* = 7 rats.

Deflating these behavior-aligned components from each animal’s CA1 activity reduced CA1- to-PFC prediction *R*^2^ by 85.0 *±* 12.4%, substantially more than deflating random dimensions of matched rank (57.6 *±* 1.3% reduction, *t* = 5.52, *P* = 0.0015, *d_z_* = 2.09, paired *t*-test, Fig. 4D). The ShaReD reduction exceeded the random reduction at each of the first 6 component counts (*P <* 0.05, Bonferroni-corrected for 6 comparisons).

ShaReD deflation reduced position decoding from *R*^2^ = 0.356 to *R*^2^ = 0.259, well beyond the reduction from random deflation of matched rank (*R*^2^ = 0.308, Fig. 4E, left). The excess reduction beyond random was significant (per-animal empirical-null tests combined by signed Liptak–Stouffer, *Z* = *−*5.73, *P* = 1*×*10*^−^*^8^, Bonferroni-corrected). Prospective choice decoding during the stem phase (the central arm of the W-track, before the choice point) was not significantly reduced by ShaReD deflation (65.2%) relative to random-deflated controls of matched rank (65.9%, n.s. after Bonferroni correction, Fig. 4E, right). The full CA1 population (67.3%) is shown for context. Position remained partly decodable from CA1 after ShaReD deflation. CA1 represents space across many cells with nonlinear place fields, so the linear behavior-aligned directions removed here capture only part of that code, and position information persists in the CA1 dimensions left untouched.

Individual ShaReD components showed distinct spatial and head-direction tuning on the W-track [15], with similar tuning across animals (Fig. 4F). The behavior-aligned components captured a large fraction of the CA1-to-PFC predictive structure during navigation while leaving signals supporting prospective choice intact. ShaReD deflation of the reverse PFC-to-CA1 communication subspace produced a similar non-significant reduction in choice decoding (Supplementary Fig. S16), so the dissociation between position and choice loss in Fig. 4E is not specific to a single direction.

## Discussion

ShaReD starts from a simple constraint. Many multi-subject experiments do not provide matched neurons, matched neural trajectories, or matched behavioral time points. They do provide paired neural and behavioral measurements within each subject. ShaReD uses that structure directly, searching for directions in behavioral feature space whose neural correlates recur across subjects while allowing each subject its own neural projection. Existing multi-subject methods put the burden of correspondence on the neural data, aligning activity across subjects through matched time points, stimuli, or conditions. ShaReD places it on the behavior instead, which provides the shared coordinate system, while neural representations are allowed to vary across subjects.

The motor cortex analysis shows how ShaReD can be used to identify shared encoding. Across macaque reaching sessions, ShaReD recovered kinematic components, each a weighted combination of hand position, distance from center, and speed, that generalized across animals and across reaching tasks. Their temporal weight profiles were smooth across lags, and the same features recurred in both tasks.

The hippocampal-prefrontal analysis shows a different use. Here a behavioral variable encoded in both regions is a potential confound rather than a target. Reduced-rank regression identifies CA1 activity patterns that predict PFC activity, but both areas encode spatial variables during navigation, and it is not clear a priori how much of this predictability can be accounted for by these behavioral variables. ShaReD can be used to remove the part of the prediction tied to specific measured behaviors. Because it fixes one behavioral projection across all animals, the same directions are deflated from every rat, and the deflation becomes a confound control matched across the whole population rather than one fit to each animal, which could capitalize on animal-specific noise. Position can already be decoded from each animal’s communication subspace on its own, so the added value here is this shared axis. Deflating these directions reduced position decoding and CA1-to-PFC prediction far more than random deflation of matched rank, while leaving prospective choice decoding indistinguishable from baseline. The same dissociation held when the prediction direction was reversed (Supplementary Fig. S16). Part of the position signal therefore resides in the shared behavior-aligned directions, though most of it lies outside of them. The activity predicting the upcoming turn, by contrast, was not captured by the recorded kinematics and survived their removal. This is what we would expect if choice-predictive activity reflects an internal variable, such as the intended turn, that is only loosely tied to the rat’s instantaneous position and head direction on the stem. The same situation arises in any communication-subspace analysis between regions that share an encoded behavioral variable, and deflating ShaReD components offers a population-matched way to remove its contribution.

A convenient mathematical property of ShaReD is the analytical elimination of the subject-specific neural projections **a***_k_* via their closed-form optima, reducing the multi-subject optimization to a single generalized eigenvalue problem over the shared behavioral space (see equation (3) in the Methods). The squared-correlation objective has two roles in this reduction. Algebraically, it enables this closed-form solution via eigendecomposition, provided no regularization terms are included. Structurally, it linearizes the way subjects’ contributions combine, so the optimum is generally located at an unmixed component rather than at a mixture of two true components whose aggregate strength across subjects is similar (Supplementary Note 1). This separates ShaReD from multiset CCA and regularized generalized CCA, which assign an independent loading vector to every block and maximize functions of pairwise correlations without a shared-loading constraint [11, 16].

The cross-animal macaque comparison with GCCA-based alignment illustrates when behavior-based alignment is preferable (Fig. 3C,D). With temporal correspondence available, behavior-mediated alignment and neural alignment produced similar cross-animal position decoding. After this correspondence was disrupted, GCCA performance degraded while ShaReD results were unchanged. Alignment-based methods remain appropriate when matched samples exist. ShaReD applies more generally when subjects share a behavioral feature space, but not necessarily a common timeline or shared trial structure.

### Limitations and practical considerations

ShaReD currently uses a stationary model based on linear correlations. This makes the components interpretable and the optimization efficient, but limits the method’s ability to capture nonlinear neural-behavioral relationships or changes in encoding over learning, context, or time. Time-varying and nonlinear extensions could address these cases at the cost of a less transparent model.

By construction, **b** captures only what subjects share. It is the combination of behavioral features whose neural correlate is most consistent across the population of individuals under the squared-correlation criterion, and it therefore reflects how the population encodes behavior rather than how any one subject does. When animals encode behavior in systematically different ways, **b** settles on the shared structure and sets aside individual idiosyncrasies. The cross-task transfer illustrates this, as components learned in one reaching task stayed predictive in the other and tracked the difficulty of the test task rather than the training task, indicating that **b** captures structure shared across reaching repertoires rather than a single task’s encoding. Individual departures from the common code are not lost but remain in the residual, which a complementary analysis could use to ask how subjects differ rather than what they share.

The method requires a shared behavioral feature space. All subjects must have the same measured behavioral variables, and missing behavioral modalities are not handled directly. Feature choice therefore matters. In the motor cortex analysis, the feature set was obtained from a sweep over candidate kinematic variables and temporal windows. In less constrained behaviors, automated behavioral feature learning, for example with autoregressive hidden Markov models or related unsupervised methods [17, 18, 19, 20, 21] may be useful, provided that the learned features remain interpretable.

The residual structure after behavioral deflation should not be read as a single clean signal. It can contain task variables that the recorded behaviors did not include, such as prospective choice [22], behaviors we did not measure such as arousal or posture, and neural activity unrelated to behavior altogether. Interpreting any residual subspace requires richer behavioral measurements and task-specific controls.

ShaReD is suited for experiments in which subjects explore overlapping behavioral repertoires without matched timing. It does not replace neural alignment when direct correspondence is available, but provides a complementary route for multi-subject analysis. It is broadly applicable as long as there is a minimal shared structure consisting of a common behavioral feature space being explored by different individuals, rather than requiring temporally aligned trajectories.

## Acknowledgements

We thank Assaf Ramot, Takaki Komiyama, Mikio Aoi, Bing Brunton, and Stefano Fusi for discussions on the method and analyses, the Miller lab for making the macaque reaching dataset publicly available, and the Jadhav lab for making the hippocampal-prefrontal dataset publicly available. This work was supported by NIH grant R01NS125298 (NINDS) and the Kavli Institute for Brain and Mind.

## Author contributions

F.H.T. and M.K.B. conceived the project and designed the method. F.H.T. implemented the algorithm, performed all analyses, and prepared the figures. F.H.T. and M.K.B. wrote the manuscript. M.K.B. supervised the work.

## Methods

### Datasets

#### Non-Human Primate Motor Cortex

The first dataset comprises Utah-array recordings of single- and multi-unit activity from primary (M1) and dorsal premotor cortex (PMd) in three macaques (C and M, both *M. mulatta*, and T, *M. fascicularis*) performing center-out (CO) and random target (RT) reaching tasks [23, 4, 24].

In the CO task, monkeys controlled a screen cursor with a planar manipulandum. Each trial began with the animal centered in the workspace. After a variable waiting period, one of eight radially arranged targets (spaced 45*^◦^* apart) appeared. Following an auditory go cue, animals had 1 s to reach the target and to hold for 0.5 s for a liquid reward. We analyzed 48 sessions from monkey C, 20 from monkey M, and 5 from monkey T. Sessions required at least 30 neurons, with CO sessions additionally requiring at least 20 trials per target direction. A fourth monkey (J) was excluded owing to low session count and unit yield (18–38 units per session).

In the RT task, monkeys made four consecutive reaches to randomly positioned targets within a 10 *×* 10 cm workspace. Each target appeared within an annulus (5 cm inner radius, 15 cm outer radius) of the previous target position to enforce consistent reach amplitudes. No explicit go cue was provided, and successive reaches were separated by brief hold (100 ms) and delay (100 ms) periods. We analyzed 15 sessions from monkey C, 6 from monkey M, and 5 from monkey T. RT sessions had no per-target trial requirement, since targets were randomly positioned.

Hand kinematic and neural data were obtained from the publicly released NWB files in DANDI dataset 000688 [24]. Cursor position and velocity were stored at 100 Hz, and spike times at their original resolution. Sessions contained 33–295 units across both tasks (Supplementary Table S3). Detailed experimental procedures are described in [4].

#### Rat Multi-Region Recordings

The second dataset comprises simultaneous hippocampal CA1 and prefrontal cortex (PFC) recordings from seven Long-Evans rats performing a spatial alternation task on a W-track maze, with continuous alternation between left and right choice arms [12]. Sessions contained 26–145 CA1 units and 21–68 PFC units from chronically implanted multi-tetrode arrays. Detailed experimental procedures are described in [12].

#### Sample Sizes and Ethics

Sample sizes (*N* = 3 monkeys, *N* = 7 rats) reflect the available data from the original studies. All animal procedures were approved by the original study institutions and are described in the respective publications.

### Data Preprocessing

#### Non-Human Primate Preprocessing

##### Neural data

Spike trains were binned at 30 Hz (33.3 ms bins). Units with mean firing rate below 1 Hz were excluded. Counts were square-root transformed for variance stabilization, smoothed with a Gaussian kernel (*σ* = 50 ms), and z-scored within each cross-validation fold in each session, following established methods for motor cortical data analysis [25, 23, 4].

##### Movement onset detection

Onset detection located the first peak in acceleration, computed as the per-sample velocity increment, exceeding 1.9 cm/s (*∼*190 cm/s^2^ at the native 100 Hz sampling rate). Onset was taken as the last preceding sample at which acceleration had fallen below half the peak value.

##### Behavioral feature extraction

The feature set comprised hand position coordinates (*x*, *y*), speed (magnitude of the velocity vector), and distance from workspace center. Feature selection was validated by a 9-configuration ablation (drop-feature, swap-direction, add-feature, and three temporal-window choices, Supplementary Fig. S10). Each variable was delay-embedded with 31 lags spanning *−*500 to +500 ms relative to neural activity at 33.3 ms resolution, yielding 124 features (4 variables *×* 31 lags) per time point. Features were grouped by kinematic variable for smoothness regularization in ShaReD. Behavioral features were z-scored within each session.

##### Quality control

Units required a mean firing rate *≥* 1 Hz. CO trials required a recorded go cue and a detected movement onset. For continuous-time data analyses, time points with hand position outside *±*10 cm of workspace center were excluded as potential tracking artifacts.

#### Rat Multi-Region Preprocessing

##### Neural data

Spike trains were binned at 100 ms, square-root transformed for variance stabilization, and smoothed with a Gaussian kernel (*σ* = 50 ms). The counts were mean-centered for the reduced-rank regression (RRR) fit, and the resulting 11-dimensional RRR latents were z-scored per cross-validation fold before ShaReD.

##### Behavioral feature extraction

The behavioral features came from four tracked variables, namely position *x*/*y* coordinates (centered, cm), 2D speed (cm/s), and instantaneous movement direction. Let *v_x_* and *v_y_* be the planar velocity components and 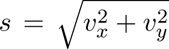 the speed. The movement direction is the angle of the velocity vector, and we represented it by its cosine and sine, (*v_x_/s, v_y_/s*), which are the components of the unit velocity vector. Splitting direction into these two components gives five features in total, and each was delay-embedded with 6 causal lags (*−*500 to 0 ms in 100 ms steps), yielding 30 features per time point.

##### Quality control

Sessions with stable recordings from both CA1 and PFC were included. Units were included based on standard spike-sorting quality metrics described in [12], with a minimum firing rate of 0.1 Hz.

### Shared Representation Discovery (ShaReD)

#### Mathematical Framework

For *K* subjects with paired neural and behavioral datasets (**X**_1_, **Y**_1_)*, . . .,* (**X***_K_,* **Y***_K_*), the neural data matrix **X***_k_ ∈* R*^Tk ×Nk^* holds *N_k_* neurons across *T_k_* time points and the behavioral data matrix **Y***_k_ ∈* R*^Tk ×B^* holds *B* behavioral features over the same time points. *T_k_* and *N_k_* vary across subjects, while behavioral feature dimensionality *B* is constant.

ShaReD identifies a shared behavioral projection **b** *∈*R*^B^* and subject-specific neural projections **a***_k_ ∈* R*^Nk^* by minimizing

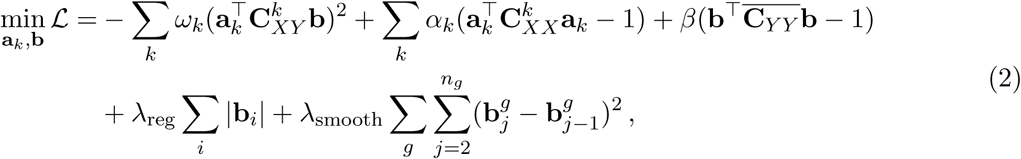

where 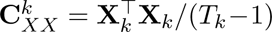 is the neural covariance, 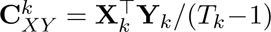 the cross-covariance, and 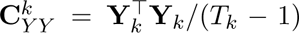 the behavioral covariance. The average behavioral covariance is 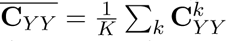. The weights *ω_k_* can scale each subject’s contribution, for instance to upweight datasets with more reliable recordings, and we set *ω_k_* = 1*/K* throughout this work, giving every subject equal influence. The L1 penalty controlled by *λ*_reg_ emphasizes features that are consistently encoded across subjects and enforces sparsity. The smoothness penalty controlled by *λ*_smooth_ applies separately to groups of temporally related features, where *g* indexes the behavioral variable type and *b^g^* denotes the projection weight for the *j*-th temporal lag of variable *g*.

#### Optimization Procedure

The optimization has two stages. The objective in equation 2 constrains each projected time series to unit variance, 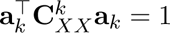 and 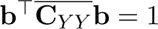, and the optimization is carried out in this constraint metric. The vectors we report and plot are rescaled to unit Euclidean norm, **â***_k_* = **a***_k_*/||**a***_k_*||_2_ and **b̂** = **b**/||_2_. The reported correlations and cosine similarities are invariant to this rescaling. The first stage of the optimization procedure solves the unregularized problem through eigende-composition. Setting partial derivatives of the Lagrangian (without the regularization terms) to zero yields an eigenvalue problem that allows us to find the shared behavioral projection **b** as proportional to the principal eigenvector of

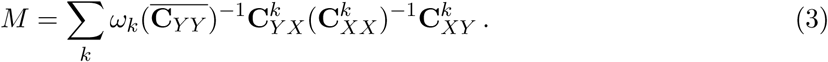

Given **b**, the neural projection for each subject is obtained in closed form as

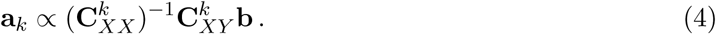

The full derivation of the optimal projection vectors is given in Supplementary Note 2.

ShaReD generalizes two-block canonical correlation analysis [9] to *K* paired neural-behavioral datasets. Classical CCA finds projections (**a**, **b**) that maximize corr(**Xa**, **Yb**) for a single (*X, Y*) pair and reduces to a generalized eigenvalue problem. ShaReD preserves the neural-behavioral pairing within each subject, ties the behavioral loading **b** across all *K* subjects while keeping the neural loadings **a***_k_* subject-specific, and sums squared rather than linear correlations to handle heterogeneous encoding strengths. Substituting each **a***_k_* with its closed-form optimum (equation (4)) removes the neural projections from the objective and reduces the joint optimization to a single eigenvalue problem over the behavioral space, the matrix *M* . ShaReD differs from multiset CCA and regularized generalized CCA [11, 16], which assign an independent loading vector to every block and maximize functions of pairwise block correlations without a shared-loading constraint.

The second stage of the optimization procedure refines this solution by gradient descent, with the L1 penalty applied via a proximal operator and the smoothness penalty entering as a smooth gradient term. Gradients with respect to neural and behavioral projections, and the smoothness gradient with boundary conditions, are given in Supplementary Note 2. At each iteration the projections are retracted onto the constraint surfaces, 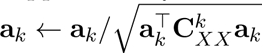 and 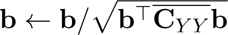, and the Lagrange multipliers are set to their unit-variance values 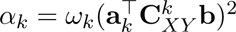 and *β* = Σ_*k*_ *α_k_*.

The retraction returns the projections to the constraint surface at each step, which keeps these multiplier values consistent with the current projections. On completion, the unit-Euclidean-norm projections **a**^*_k_* and **b**^^^ are orthogonalized against previously found components via Gram–Schmidt and renormalized to unit Euclidean norm.

For numerical stability, the neural covariances 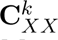 and the pooled behavioral covariance 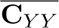 are regularized by adding *ɛ*_cov_*I* where *ɛ*_cov_ = 10*^−^*^6^. Matrix inversions in the eigendecomposition step use a ridge-regularized inverse with *ɛ*_inv_ = 10*^−^*^10^. Gradient descent uses learning rates initialized at 0.01 with exponential decay (factor 0.99 every 100 iterations) and runs for 1,000 iterations. Stage 1 solves the unregularized problem exactly, so it lands close to the optimum of the full regularized objective (Supplementary Fig. S12B). Stage 2 therefore amounts to light refinement, and the schedule above suffices.

#### Multi-Component Extraction and Deflation

Multiple components are extracted via deflation [26]. After identifying component *i* with behavioral projection **b**^^(^*^i^*^)^ and neural projections 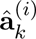, the data are deflated as 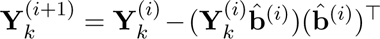 and 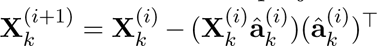. The procedure continues until the desired number of components is extracted or the remaining variance becomes negligible.

#### Implementation Details

##### Preprocessing

**X***_k_* and **Y***_k_* were z-scored within session before being passed to ShaReD, as described in Data Preprocessing. The behavioral matrix **Y***_k_* was then ZCA-whitened per session and per fold. Let *V* Λ*V ^⊤^* be the eigendecomposition of the training-fold sample covariance with eigenvalues *λ_i_*, and let the whitening transform be *W* = *V* diag max(*λ_i_, ɛ*_zca_) *^−^*^1^*^/^*^2^*V ^⊤^*, with eigen-value floor *ɛ*_zca_ = 10*^−^*^8^ for numerical stability. *W* was computed on the training fold and applied unchanged to the matched test fold. For the primate motor cortex data the neural matrix **X***_k_* was whitened the same way, whereas for the rat multi-region data the 11-dimensional RRR latents were passed in with z-scoring only, since they are already a low-rank projection of CA1 activity and well conditioned by construction. Whitening of **Y***_k_* leaves the pooled 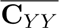 close to the identity, so *M* in equation (3) is close to symmetric and a standard symmetric eigensolver is well behaved in practice. Whitening of **X***_k_* is not required for correctness, since the closed-form solution uses the explicit inverse 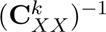. For the primate analyses it was applied as a numerical conditioning step.

##### Hyperparameter selection

Regularization parameters were selected by nested 10-fold cross-validation. Within each outer fold the model was fit on a sub-training partition, hyperparameters were chosen on a validation block taken from the adjacent outer fold, and the held-out test block gave the unbiased correlation. For the primate motor cortex analyses, *λ*_reg_ and *λ*_smooth_ were each swept over *{*0.0, 0.01, 0.02, 0.05, 0.1, 0.2, 0.3, 0.5, 1.0, 2.0, 5.0, 10.0*}*, choosing values that maximized held-out correlation while preserving component interpretability. Standard primate analyses used *λ*_reg_ = 0.01 and *λ*_smooth_ = 0.01, and the rat multi-region analysis used *λ*_reg_ = 0.001 and *λ*_smooth_ = 0.0.

#### Synthetic Data Validation

Synthetic datasets were generated from a linear latent factor model, 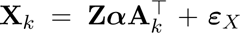 and **Y***_k_* = **Z*β*B***^⊤^* + ***ε****_Y_*, with orthonormal subject-specific neural loadings **A***_k_*, shared orthonormal behavioral loadings **B**, orthonormalized latent factors **Z**, Gaussian noise (*σ* = 1.0), and target component correlations set by symmetric diagonal scaling ***α*** = ***β***. Data were z-scored and whitened per subject before fitting. In each analysis below, recovered components were matched to ground truth by the Hungarian algorithm [10] and recovery was reported as the absolute cosine similarity between estimated and ground-truth projections, averaged over 100 random-seed repetitions. Full generative model, correlation specification, preprocessing, and per-experiment parameters are in Supplementary Note 3.

##### Component recovery across signal strengths

ShaReD was fit to datasets with 14 ground-truth components and target correlations from 0.05 to 0.70 across 6 subjects, using the standard validation configuration (Table S1, Fig. 1B). Test performance was additionally reported as the absolute Pearson correlation *|r|* between projected neural and behavioral data on held-out samples, matching the sign-invariant cosine similarity used for component recovery.

##### Sensitivity to sample size, noise, and number of subjects

Three one-at-a-time sweeps, shown in Supplementary Fig. S3, varied training set size, noise level *σ*, and number of subjects *K* around a fixed baseline (*T*_train_ = 2000, *K* = 6, *σ* = 1.0).

##### Recovery of components with different sharing structure

Datasets were generated with five components across seven subjects, comprising one global component, two subgroup components splitting the population, and two pair components matched in aggregate strength but differing in which two subjects share them. All components targeted information about the sharing structure (Fig. 1C). *ρ̄* = 0.60. ShaReD was fit without

##### Cross-subject pseudopopulation decoding

Behavioral data were projected onto each ShaReD component direction and binarized by median split. Pseudopopulations were constructed by concatenating neural activity across subjects, and logistic regression classifiers were trained on these pseudopopulations to predict the binary label. Three baselines were used, random projection directions in behavioral space, individual behavioral feature dimensions in the whitened frame, and a shuffled-label control (Fig. 1D).

##### Objective function comparison

Optimization landscapes were analyzed for linear (*p* = 1), squared (*p* = 2), and quartic (*p* = 4) correlation objectives, both analytically and through numerical sweeps over mixing angle and cross-subject correlation heterogeneity. The full analysis is described in Supplementary Note 1.

##### Component identifiability versus single-subject CCA

CCA cannot reliably separate two components whose canonical correlations are close, since the underlying eigenvalue problem becomes degenerate as the correlations converge. ShaReD inherits this limitation through its closed-form eigendecomposition but can mitigate it by pooling samples across subjects. We characterized both effects on two-component datasets with one component anchored at correlation *ρ*_1_ and the second at *ρ*_1_ + Δ, comparing ShaReD (*K* = 6) against single-subject CCA (*K* = 1) under two baselines, matched per-subject *T* (CCA sees one subject’s samples) and matched total *T* (CCA sees the combined sample count of all subjects in a single subject). Each method was evaluated across anchor strength and sample size (Supplementary Fig. S2).

### Primate Analyses

#### Within-Task Application

For each task, ShaReD was fit jointly across all CO or RT sessions, with each session contributing its own **X***_k_* and **Y***_k_* and the algorithm returning one shared behavioral projection **b** together with session-specific neural projections **a***_k_*. We restricted all reported analyses to the leading 20 components. We used block-randomized 10-fold cross-validation. Each session was partitioned into 20 contiguous blocks of approximately equal length, and whole blocks were assigned to the 10 folds at random. A 30-sample gap (*±*15 lags at 33.3 ms resolution, totaling 1 s) was left between consecutive blocks to prevent temporal leakage from the delay-embedded behavioral features. Whitening was applied per fold as described in ShaReD Implementation Details. Component significance was assessed against a refit-shuffle null. The behavioral matrix was independently shuffled within each session’s training fold, ShaReD was refit by closed-form eigendecomposition without regularization at the maximum number of components the data support, set by the smaller of the behavioral rank and the smallest per-session neural rank, and test correlations were recomputed on the unshuffled test fold. The per-component test statistic was the RMS cross-validated neural-behavioral correlation across sessions. The null was formed by pooling the shuffled statistics across the 10 folds, so the 100 shuffles produced 1,000 draws. Per-component *p*-values were corrected across components by the Benjamini–Hochberg false discovery rate, which yields per-component *q*-values (*p*_FDR_), and the reported significant set is the leading contiguous run with *q <* 0.05.

#### Leave-One-Animal-Out (LOAO)

For cross-animal generalization, ShaReD was trained on all sessions from *N −* 1 animals to identify the shared **b**, which was then frozen. For each session of the held-out animal, 10-fold cross-validation was applied. Within each fold, the training portion of the session was used to compute the session-specific neural projection **a***_k_* in closed form (equation (4)), and the Pearson correlation between **X***_k_***a***_k_* and **Y***_k_***b** was evaluated on the held-out test portion. Reported correlations are averages across the 10 folds. Chance levels were obtained by shuffling the held-out animal’s neural matrix instead of the behavioral matrix, preserving the frozen behavioral projection while breaking within-animal neural-behavioral correspondence.

#### Cross-Task Generalization

ShaReD was trained on all sessions from one task (e.g., CO) to identify the shared **b**, which was then frozen. Each session of the other task was evaluated by 10-fold cross-validation as in the LOAO analysis, with the session-specific neural projection **a***_k_* fit on the training portion in closed form (equation (4)) and the correlation evaluated on the held-out test portion.

#### Comparison with GCCA

ShaReD was compared against generalized canonical correlation analysis (GCCA) using the MAX-VAR formulation [11] in two settings, pairwise cross-session decoding and leave-one-animal-out generalization.

##### Pairwise cross-session decoding

Trial epochs spanned *−*50 to 400 ms relative to movement onset. For pairwise analysis, GCCA was used to align neural spaces between each pair of sessions.

Each session’s neural data was first reduced to 10 principal components, then GCCA was fit to find a shared representation space *G* and session-specific weights by minimizing ∑_*s*_||*G* − **X***_s_W_s_*||^2^ subject to *G^⊤^G* = *I*. The alignment was fit on the training-fold PCA data of both sessions, matching the ridge decoder’s fold split. Because GCCA is unsupervised with respect to the decoding target, this scoping does not introduce position leakage. A ridge regression decoder (*α* = 1.0) was trained on GCCA-projected training data from one session and tested on GCCA-projected test data from the other session, predicting 2D position coordinates (*x, y*). Reported *R*^2^ is averaged across 6 trial-stratified folds.

In pairwise mode, ShaReD was fit on the same 10-component PCA representation of both sessions jointly to identify shared behavioral projections and session-specific neural projections. A ridge regression decoder trained on ShaReD-projected data from one session was tested on ShaReD-projected data from the other, with *R*^2^ averaged across the same 6 trial-stratified folds.

##### Leave-one-animal-out (LOAO) decoding

For LOAO analysis, GCCA was fit on the training-fold PCA-reduced neural data from all sessions, including those from the held-out animal. Because GCCA’s alignment is unsupervised with respect to position, including the held-out animal’s neural data in the alignment does not leak the decoding target, so holdout was applied only at the decoder. A ridge regression decoder trained on pooled GCCA-projected data from training animals was tested on the held-out animal’s sessions, with *R*^2^ averaged across 6 trial-stratified folds within held-out sessions.

For ShaReD, the neural data were reduced to the same 10 principal components, and **b** was fit on training animals only and then frozen. For each held-out session, **a***_k_* was fit on training folds to maximize correlation with the frozen **b**, and a ridge regression decoder trained on pooled ShaReD-projected training data was tested on the projected test fold of the held-out session.

##### Temporally aligned versus shuffled data

In the temporally aligned condition, epoch sequences matched across sessions. In the shuffled condition, the target labels of one session were randomly permuted, re-pairing trials across sessions. The same shuffle protocol applied to GCCA and ShaReD. For pairwise analysis, the neural activity, the behavioral feature matrix used for alignment, and the hand position matrices used as the decoding target were jointly permuted with a single random seed while the other session was left intact, preserving within-session correspondence while breaking cross-session temporal alignment. For LOAO analysis, the shuffle was applied to held-out sessions only, with each session shuffled jointly using its own seed, and training data were left intact. Because GCCA relies on neural-to-neural alignment across sessions, shuffling disrupts its ability to align or project held-out data. ShaReD learns projections from within-session neural-behavioral correspondence and is therefore unaffected by cross-session misalignment.

### Rat Multi-Region Analyses

#### Communication Subspace Extraction (RRR)

For each animal, reduced-rank regression was fit to predict PFC from CA1 activity. Training data were mean-centered (not z-scored), following [14, 12]. A 5-fold cross-validated rank sweep over *r ∈ {*1*, . . .,* 30*}* gave per-animal one-standard-error (1-SE) cutoffs (SE across folds) with median *r* = 11 and range 5–15 across the 7 animals (Supplementary Fig. S14), and we used this median rank, *r* = 11, for all animals. Ridge regularization was set to 10*^−^*^10^ (near full-rank solution).

Cross-validation used 5 folds split on unique trial IDs so that all timepoints from a trial remained in the same fold. The rank-*r* fit identifies an *r*-dimensional subspace of CA1 activity that best predicts PFC. Let *U_k_ ∈* R*^N^*^CA1^*^,k ×r^* be an orthonormal basis for this subspace, the right-singular vectors of the fit. Projecting each animal’s CA1 activity onto this basis gives the *r*-dimensional latents *Z_k_* = (**X**_CA1_,_*k*_ − *X̄_k_*)*U_k_*, which are the inputs to ShaReD.

#### ShaReD on Communication-Subspace Latents

ShaReD was fit to the RRR-derived latents across animals, extracting all 11 shared components of the CA1-to-PFC communication subspace by 5-fold cross-validation aligned with the RRR folds, with *λ*_reg_ = 0.001 and *λ*_smooth_ = 0.0. Preprocessing followed ShaReD Implementation Details. Component significance used the refit-shuffle null of Within-Task Application, with one difference. The behavioral projection **b** was refit jointly across all 7 animals on each shuffle while each animal’s **a***_k_* was obtained in closed form, giving a single across-animal RMS statistic and one *p*-value per component rather than per-animal *p*-values combined afterward. Pooling the 100 shuffles across the 5 folds gave a null of 500 draws, corrected across components by Benjamini–Hochberg FDR and reported as *q*-values (main text and Supplementary Fig. S15).

#### Deflation in the Communication Subspace

ShaReD components were deflated from CA1 activity by orthogonal projection. The ShaReD neural projections live in the z-scored latent frame, so removing them from raw CA1 activity requires mapping them back to CA1 space. Let *Q_A_* = QR(*A*) be an orthonormal basis for the *c* ShaReD neural projections in that frame. Undoing the per-dimension z-scoring of animal *k*’s latents (*s_k_* = diag(*σ_k_*)) and re-embedding through the communication-subspace basis *U_k_* carries this basis into CA1 space, where 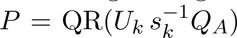 is the resulting orthonormal projector. CA1 activity was deflated as 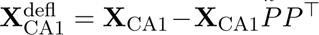. For the CA1*→*PFC re-evaluation the same components were removed from the z-scored latents and mapped back before the frozen RRR model. Random deflation controls used 100 matched-rank random orthonormal projections from the communication subspace span(*U_k_*).

#### Position and Choice Decoding

Decoders were evaluated on outbound trials only, since inbound trials lack a decision component. Task phase (stem versus choice) was inferred per timepoint from the rat’s position relative to the T-junction of the W-track [12].

*Position decoding* predicted the (*x, y*) coordinate at each timepoint via ridge regression (*α* = 100.0) with z-scored inputs and targets. The metric was *R*^2^ averaged across the two output dimensions. Both stem and choice phases were used to capture the full W-track spatial variability.

Neural features were clipped at *±*5 SD after z-scoring to prevent rare firing-rate outliers from dominating the ridge solution. Latent-space features (projections and residuals) were well conditioned and require no clipping.

*Choice decoding* predicted upcoming turn direction (left versus right, binary) via logistic regression (inverse L2 regularization strength *C* = 1.0, balanced class weights), evaluated by balanced accuracy. Stem-phase data were used with causal-only delay embedding (*−*5 to 0 lags, i.e. 6 lags spanning *−*500 to 0 ms at the 100 ms bin resolution), so classifiers predict the upcoming turn from pre-decision neural activity without leakage from post-decision trajectory features. Chance controls used 100 label permutations.

*Cross-validation for decoding.* Both decoders used the same 5-fold splits as the RRR and ShaReD stages. Within each fold, we compared the full CA1 population (baseline), ShaReD-deflated CA1, and random-deflated CA1. Statistical procedures for these comparisons are described in Statistical Analysis.

### Statistical Analysis

#### Performance Metrics

Component recovery on synthetic data was scored as the absolute cosine similarity between true and estimated projection vectors, with the Hungarian algorithm [10] used to match recovered components to ground truth and account for ordering ambiguity. Held-out performance was reported as Pearson correlation (*r*) or coefficient of determination (*R*^2^), depending on the analysis.

#### Hypothesis Testing

For the progressive deflation sweep (Fig. 4D), paired *t*-tests across the 7 rats compared deflation conditions at each rank. For the decoding dissociation (Fig. 4E), per-animal *p*-values were computed against an empirical null of matched-rank random deflations and combined across the 7 animals by the signed Liptak–Stouffer procedure, with the combined *p*-value taken from the normal approximation of the Stouffer *Z* statistic. The full (intact-CA1) condition in Fig. 4E was included for visual context. Only the ShaReD-deflated versus random-deflated contrast was formally tested. Mann–Whitney *U* tests compared per-component correlations between tasks (e.g., CO vs RT, Fig. 2B) and behavioral variability across tasks. For the aligned-versus-shuffled comparison of cross-animal decoding (Fig. 3C,D), GCCA’s change in *R*^2^ was tested with a clustered sign-flip permutation test (10,000 permutations) rather than a paired test, because the units are not independent. In the pairwise analysis the 1,300 cross-animal pairs are built from 73 sessions, so each session recurs across many pairs, and treating the pairs as independent would overstate the significance. The statistic was the mean aligned-minus-shuffled *R*^2^, and the null was generated by randomly flipping the sign of each session cluster’s contribution. Clusters were sessions in both designs. In the pairwise analysis (C) each cross-animal pair carries two session memberships and was flipped by the product of its two sessions’ signs, and in the leave-one-animal-out analysis (D) each observation was a single session. Clustering was done at the session level. ShaReD’s fit runs through per-session covariances, which are invariant to within-session reordering, leaving its solution unchanged under the same shuffle. ShaReD is therefore shown for reference rather than tested.

#### Multiple Comparison Correction

Multiple comparison corrections were applied per analysis. The GCCA comparisons (Fig. 3C,D), the progressive deflation sweep (Fig. 4D, 6 rank comparisons), the decoding dissociation (Fig. 4E, 2 comparisons), and the reversed-direction analysis (Supplementary Fig. S16, 2 comparisons) each used Bonferroni correction within the panel. Per-component significance tests used the Benjamini– Hochberg false discovery rate at *q <* 0.05.

#### Error Estimation

Error bars and shaded regions represent standard deviation or standard error of the mean as indicated in each figure caption. Cross-validation provided unbiased estimates of generalization performance.

#### Significance Thresholds

Unless otherwise stated, statistical significance was set at *P <* 0.05. Effect sizes are reported where applicable.

### Software and Computational Resources

#### Implementation

ShaReD targets Python *≥* 3.10. The implementation will be released publicly under the MIT license upon publication. Declared dependency floors in pyproject.toml are numpy *≥*1.20, scipy *≥*1.7, scikit-learn *≥*1.0, matplotlib *≥*3.3, pandas *≥*1.3, seaborn *≥*0.11, and h5py *≥*3.0. The figures in this paper were produced with Python 3.12 and the dependency versions pinned in the repository environment file.

#### Computational Requirements

Per component, the eigendecomposition step has complexity 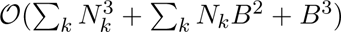, where *N_k_* is the neural dimensionality for subject *k*, *B* is the behavioral dimensionality, and *K* is the number of subjects. The first term reflects neural covariance matrix inverses for each subject, the second the per-subject contributions to the behavioral-space matrix *M*, and the third the final eigendecomposition. Covariance matrices are rebuilt on the deflated data at each component, so extracting *C* components multiplies the eigendecomposition cost by *C*. ZCA whitening via eigendecomposition of the sample covariance adds an 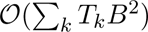 preprocessing cost, which can dominate when *T_k_* is large. With similar neural dimensions *N* across subjects, the per-component cost simplifies to *O*(*KN* ^3^ + *KNB*^2^ + *B*^3^). For typical applications (100–500 neural dimensions, 100–200 behavioral features, 3–10 subjects), computation takes seconds to minutes on standard hardware.

#### Reproducibility

Random seeds were fixed (seed=100) for stochastic procedures including data shuffling and cross-validation splits. The code includes unit tests (pytest) verifying mathematical correctness against known solutions. All figures are reproducible from the included analysis scripts.

#### Data Availability

Primate motor cortex data are from [4, 23, 24] (DANDI DOI 10.48324/dandi.000688/0.250122.1735) and rat navigation data are from [12] (DANDI archive, DOI 10.48324/dandi.000978).

