## Supplement for "Shared Representation Discovery for Multi-Subject Neural Data Analysis"

### Supplementary Note 1: Mathematical Foundation of the Squared Correlation Objective

A linear correlation objective applied to multi-subject neural data can prefer a mixture of two true components over either component on its own. This occurs when the two neural-behavioral relationships have similar aggregate correlation strength across subjects. We place ShaReD's squared objective within a family that raises the per-subject correlations to a power $p$, and we show that the squared objective sits at the threshold that separates objectives preferring a mixture from those preferring the unmixed components. The argument reduces each candidate solution to a one-dimensional landscape over the mixing between two components and reads off, from the power alone, where each objective places its maximum.

This failure mode is specific to the multi-subject setting. In single-subject CCA, mixing arises only as a rotational degeneracy at an exact tie between two component correlations, and any perturbation breaks it. ShaReD inherits this exact-tie degeneracy, but as soon as the two correlations differ slightly, pooling across subjects resolves the two components more reliably than any single subject can (Supplementary Fig. S2). The mixing analyzed here is different. It occurs when individual subjects have well-separated component strengths, yet the way the objective pools them across subjects creates a preference for a mixed direction over either unmixed component.

**Per-subject reduction.** Consider $K$ subjects with paired neural and behavioral data, with each column of $\mathbf{X}_k$ and $\mathbf{Y}_k$ mean-centered, and let $\rho_{1,k}$ and $\rho_{2,k}$ denote the correlations achievable by two underlying neural-behavioral components for subject $k$. We parameterize a candidate solution by a single mixing angle $\theta$ on the behavioral side and take the neural projection at its optimum for each subject. The candidate behavioral projection $\tilde{b}(\theta)$ is a combination of the true orthogonal components $b_1^*$ and $b_2^*$,

$$\tilde{b}(\theta) = \cos(\theta)\, b_1^* + \sin(\theta)\, b_2^* \,. \tag{5}$$

For a fixed $\tilde{b}(\theta)$, the correlation that subject $k$ achieves with a neural projection $a$ is

$$\rho_k(a) = \frac{a^\top \mathbf{C}_{XY}^k \tilde{b}}{\sqrt{\left(a^\top \mathbf{C}_{XX}^k a\right)\left(\tilde{b}^\top \mathbf{C}_{YY}^k \tilde{b}\right)}} \,. \tag{6}$$

The behavioral term is fixed, so maximizing $\rho_k(a)$ over $a$ is a linear regression of $\mathbf{Y}_k\tilde{b}$ on $\mathbf{X}_k$, with solution

$$\tilde{a}_k(\theta) \propto (\mathbf{C}_{XX}^k)^{-1}\mathbf{C}_{XY}^k \tilde{b}(\theta) \,. \tag{7}$$

The overall scale of $\tilde{a}_k$ cancels in $\rho_k$, so we use this projection as written. Write $\mathbf{M}_k = \mathbf{C}_{YX}^k(\mathbf{C}_{XX}^k)^{-1}\mathbf{C}_{XY}^k$. Substituting $\tilde{a}_k$, the numerator and the neural-side variance reduce to the same quadratic form in $\tilde{b}$,

$$\tilde{a}_k^\top \mathbf{C}_{XY}^k \tilde{b} = \tilde{b}^\top \mathbf{C}_{YX}^k (\mathbf{C}_{XX}^k)^{-1} \mathbf{C}_{XY}^k \tilde{b} = \tilde{b}^\top \mathbf{M}_k \tilde{b} \,, \tag{8}$$

$$\tilde{a}_k^\top \mathbf{C}_{XX}^k \tilde{a}_k = \tilde{b}^\top \mathbf{C}_{YX}^k (\mathbf{C}_{XX}^k)^{-1} \mathbf{C}_{XX}^k (\mathbf{C}_{XX}^k)^{-1} \mathbf{C}_{XY}^k \tilde{b} = \tilde{b}^\top \mathbf{M}_k \tilde{b} \,. \tag{9}$$

Computing $\rho_k^2(\theta)$, the squared numerator $(\tilde{b}^\top \mathbf{M}_k \tilde{b})^2$ and the neural-side variance $\tilde{b}^\top \mathbf{M}_k \tilde{b}$ share a factor, so one cancels and the squared correlation reduces to a ratio of two quadratic forms in $\tilde{b}$,

$$\rho_k^2(\theta) = \frac{\tilde{b}^\top \mathbf{M}_k \tilde{b}}{\tilde{b}^\top \mathbf{C}_{YY}^k \, \tilde{b}} \,. \tag{10}$$

The matrix $\mathbf{M}_k$ carries the canonical structure, and the denominator is handled by whitening. ShaReD operates on behaviorally whitened inputs, meaning each $\mathbf{Y}_k$ is transformed so that $\mathbf{C}_{YY}^k = \mathbf{I}$ (see Methods). The squared canonical correlations are the eigenvalues of $(\mathbf{C}_{YY}^k)^{-1}\mathbf{M}_k$, which under whitening is just $\mathbf{M}_k$, so the eigenvalues of $\mathbf{M}_k$ are the squared canonical correlations, with the behavioral canonical directions as eigenvectors. Under the generative model of Supplementary Note 3, with orthonormal loadings and orthogonal latent factors, these directions are the components $b_1^*$ and $b_2^*$, and

$$\mathbf{M}_k = \rho_{1,k}^2 \, b_1^* b_1^{*\top} + \rho_{2,k}^2 \, b_2^* b_2^{*\top} \,. \tag{11}$$

With $\mathbf{C}_{YY}^k = \mathbf{I}$ the denominator is $\tilde{b}^\top \tilde{b} = \cos^2\theta + \sin^2\theta = 1$, independent of $\theta$. Expanding the numerator over the two components,

$$\tilde{b}^\top \mathbf{M}_k \tilde{b} = \cos^2\theta \left(b_1^{*\top} \mathbf{M}_k b_1^*\right) + \sin^2\theta \left(b_2^{*\top} \mathbf{M}_k b_2^*\right) + 2\cos\theta\sin\theta \left(b_1^{*\top} \mathbf{M}_k b_2^*\right) . \tag{12}$$

The diagonal terms are $b_j^{*\top} \mathbf{M}_k b_j^* = \rho_{j,k}^2$, and the cross term vanishes, $b_1^{*\top} \mathbf{M}_k b_2^* = 0$, because $b_1^*$ and $b_2^*$ are orthonormal eigenvectors of $\mathbf{M}_k$. The would-be mixing term is therefore absent, and

$$\rho_k^2(\theta) = \cos^2(\theta)\, \rho_{1,k}^2 + \sin^2(\theta)\, \rho_{2,k}^2 \,. \tag{13}$$

This neural projection maximizes the correlation $\rho_k$ for subject $k$, and the same projection is optimal at every power $p$. Each subject's neural projection enters the objective only through that subject's own correlation, and raising a correlation to any positive power is monotonic, so the projection that maximizes $\rho_k$ also maximizes $\rho_k^p$ for every $p$. The power changes how the subjects are combined, not the projection each subject contributes. We therefore determine the neural projections once, using the same $\rho_k(\theta)$ for every power, and study the resulting function of the behavioral mixing.

**The objective family.** Substituting $u = \cos^2(\theta)$ gives $u \in [0, 1]$, with $u = 1$ at the first unmixed component ($\theta = 0°$), $u = 0$ at the second ($\theta = 90°$), and $u = 0.5$ at maximal mixing ($\theta = 45°$). The achievable squared correlation is $\rho_k^2(u) = u\, \rho_{1,k}^2 + (1-u)\, \rho_{2,k}^2$. We collect the subjects into an objective that sums their correlations raised to a power $p$, weighted by $\omega_k$,

$$L_p(u) = \sum_{k=1}^{K} \omega_k \, \rho_k^p(u) = \sum_{k=1}^{K} \omega_k \, \ell_k(u)^{p/2}, \qquad \ell_k(u) = u\, \rho_{1,k}^2 + (1-u)\, \rho_{2,k}^2 \,. \tag{14}$$

We use equal weights $\omega_k = 1/K$ throughout, so the weights sum to one and match the main objective (Eq. 1), and the two-subject examples below take $\omega_k = 1/2$. The qualitative conclusions for $p \geq 2$ hold for any strictly positive weights.

**Curvature sets the optimum.** We now read off, from the power $p$ alone, whether $L_p$ prefers a mixed direction or an unmixed component. Each subject contributes $\ell_k(u)^{p/2}$ to $L_p$, where $\ell_k(u)$ is affine in $u$ and non-negative. Because $\ell_k$ is a straight line in $u$, the curvature of this contribution comes entirely from the exponent. Differentiating twice,

$$\frac{d^2}{du^2}\, \ell_k(u)^{p/2} = \frac{p}{2}\left(\frac{p}{2} - 1\right) \ell_k(u)^{\,p/2-2} \, \left(\rho_{1,k}^2 - \rho_{2,k}^2\right)^2 , \tag{15}$$

and the sign is that of $\frac{p}{2}(\frac{p}{2}-1)$, since the remaining factors are non-negative. For a subject whose two components differ in strength, the contribution is strictly concave for $0 < p < 2$, linear for $p = 2$, and strictly convex for $p > 2$. The objective $L_p$ inherits this curvature as a non-negative sum.

The curvature determines where the maximum sits. Recall that the endpoints $u = 0$ and $u = 1$ are the unmixed components and interior $u$ is a mixed direction. A concave contribution is dome-shaped, so it can peak in the interior, and for $0 < p < 2$ the objective can prefer a mixed direction. A linear contribution is largest at an endpoint, so for $p = 2$ the objective is maximized by an unmixed component, unless its slope vanishes, in which case any mixture is equally good. A convex contribution is bowl-shaped, lowest in the middle and largest at its endpoints, so for $p > 2$ the objective is again maximized by an unmixed component, and strictly so. The squared objective corresponds to the threshold $p = 2$, the smallest power at which mixing is never preferred and the single power at which each subject's contribution is linear in the mixing coordinate.

**Symmetric and asymmetric strengths.** Write $A = \sum_k \omega_k \rho_{1,k}^2$ and $B = \sum_k \omega_k \rho_{2,k}^2$ for the aggregate strengths of the two components. At $p = 2$ the objective is exactly $L_2(u) = B + u(A - B)$, maximized at the unmixed component with the larger aggregate strength, or flat when $A = B$. Consider the symmetric case of two equally weighted subjects with $(\rho_{1,1}, \rho_{2,1}) = (0.9, 0.5)$ and $(\rho_{1,2}, \rho_{2,2}) = (0.5, 0.9)$, so that $A = B$. Here $L_2$ is flat (Fig. S1A, gray). The linear objective $L_1(u) = \frac{1}{2}\sqrt{0.81u + 0.25(1-u)} + \frac{1}{2}\sqrt{0.25u + 0.81(1-u)}$ is strictly concave and symmetric about $u = 0.5$, so it peaks at maximal mixing (Fig. S1A, blue). The quartic objective is convex and peaks at the unmixed endpoints, where $L_4(0) = L_4(1) = \frac{1}{2}(0.81^2 + 0.25^2) \approx 0.36$ exceeds the mixed value $L_4(0.5) \approx 0.28$ (Fig. S1A, red). When the aggregates differ, as for Subject 1 $(0.85, 0.5)$ and Subject 2 $(0.5, 0.8)$ with $A \neq B$, the squared and quartic objectives both select the stronger component while the linear objective still prefers a mixture (Fig. S1B). Panel C sweeps the separation incentive against the correlation difference $|\Delta\rho|$ over $[0, 0.4]$ for both cases. The symmetric case fixes the mean correlation at $\bar{\rho} = 0.7$ and takes $(\bar{\rho} + |\Delta\rho|/2,\ \bar{\rho} - |\Delta\rho|/2)$ for Subject 1 with the mirror for Subject 2. The asymmetric case centers the two subjects at different bases, 0.675 and 0.65, with unequal gaps, taking $(0.675 + 0.875\,|\Delta\rho|/2,\ 0.675 - 0.875\,|\Delta\rho|/2)$ for Subject 1 and $(0.65 - 0.75\,|\Delta\rho|/2,\ 0.65 + 0.75\,|\Delta\rho|/2)$ for Subject 2, the bases and gap factors set so that $|\Delta\rho| = 0.4$ reproduces the panel-B correlations $(0.85, 0.5)$ and $(0.5, 0.8)$. In each case the incentive is evaluated in closed form as $L_p(u{=}1) - L_p(u{=}0.5)$ for $p = 1, 2, 4$, which is the caption's $L(0°) - L(45°)$ under $u = \cos^2\theta$.

At $p = 2$ the objective also admits a closed-form solution through a generalized eigenvalue decomposition (equation (33) in Supplementary Note 2 and equation (3) in the Methods), which lets ShaReD (without regularization) bypass non-convex optimization and return a solution that cannot prefer a mixed alternative.

**End-to-end recovery.** The landscape analysis predicts where each objective places its optimum, and the fitted estimator follows it. On synthetic data with the symmetric structure above, ShaReD recovers the two ground-truth behavioral loadings at $p = 4$, returns their mixture at $p = 1$, and at $p = 2$, where the flat landscape leaves the eigendecomposition degenerate, returns an arbitrary direction in the plane spanned by $b_1^*$ and $b_2^*$ that scatters between the two (Fig. S1D). We score recovery by the alignment of each recovered loading with the true loadings rather than by the achieved correlation, which separates the identity of the recovered component from the attenuation that a shared noisy behavioral readout imposes on every correlation. The $p = 2$ eigendecomposition supplies the same initialization at every power, and all three fits are then refined by gradient descent on their own objectives. Because the $p = 2$ landscape is flat, those steps leave its fit at the arbitrary eigensolution, while the $p = 1$ and $p = 4$ fits descend to a mixture and to the separated components.

**Higher powers.** Increasing the power above two sharpens the separation incentive, but with diminishing returns and a cost. A larger $p$ amplifies the strongest per-subject correlations relative to the weakest, so the sum $L_p$ leans ever more heavily on the best-aligned subjects. As $p$ grows the objective is driven by the single subject that attains the highest correlation at a given $u$, and the statistical power gained by pooling across the rest is progressively discarded.

**Scope.** The analysis applies to the extraction of a single component under orthogonal latent factors and orthonormal loadings. Whether the squared objective continues to avoid mixing after deflation depends on how closely the deflated data preserve the remaining components' orthogonal structure. We do not prove this extension formally. Empirically, we observe clean recovery across deflation steps in both synthetic (Fig. 1C-D) and biological (Fig. S9) data.

ShaReD uses the squared objective by default, which performs reliably across the datasets analyzed in this work. The closed-form eigendecomposition is computationally efficient and is guaranteed not to prefer mixed solutions. The implementation supports gradient descent refinement with a higher-power objective, initialized from the squared solution, but this was not necessary for analyzing the biological datasets presented here.

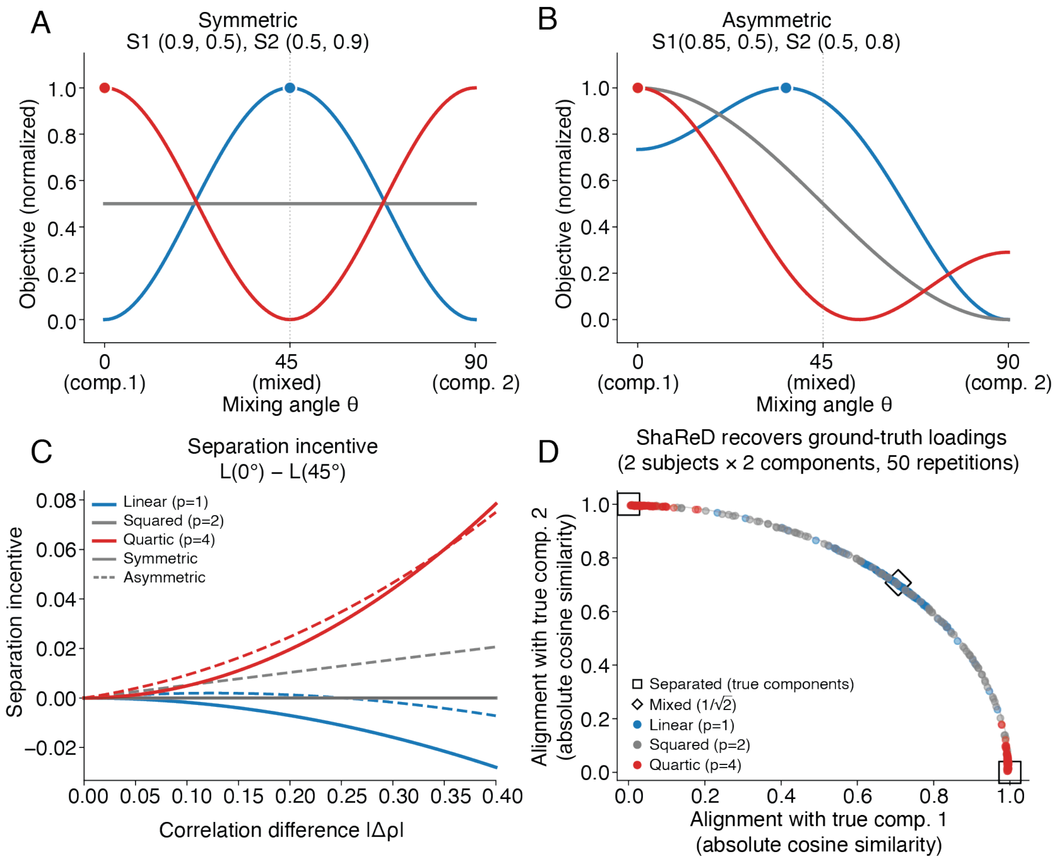


Figure S1: **Objective function landscapes under component mixing. A, B)** Objective values plotted against mixing angle $\theta$, where $0°$ and $90°$ are the unmixed true components and $45°$ is maximal mixture. Each curve is min-max normalized to $[0, 1]$ across the mixing angle, and the squared objective, which is constant in the symmetric case, is drawn at 0.5. Dots indicate global maxima. **A)** Symmetric case. Subject 1 correlations $(\rho_1, \rho_2) = (0.9, 0.5)$, Subject 2 $(0.5, 0.9)$. The linear objective (blue) peaks at the mixed solution ($45°$). The squared objective (gray) is constant, providing no gradient and no preference for mixing. The quartic objective (red) peaks at the unmixed boundaries. **B)** Asymmetric case. Subject 1 $(0.85, 0.5)$, Subject 2 $(0.5, 0.8)$. The linear objective still prefers a mixed solution. The squared and quartic objectives both identify the first unmixed component ($0°$) as optimal. **C)** Separation incentive, defined as $L(0°) - L(45°)$, as a function of the correlation difference $|\Delta\rho|$ between components, for the symmetric (solid lines, panel A) and asymmetric (dashed lines, panel B) cases. Positive values indicate a preference for unmixed solutions. The linear objective (blue) has negative separation incentive and prefers mixing in both cases. The squared objective (gray) is neutral in the symmetric case and positive in the asymmetric case, where the unequal aggregate strengths $A$ and $B$ give it a separation incentive of $(A - B)/2$. The quartic objective (red) provides a stronger separation incentive as the correlation difference grows. **D)** Recovery of the ground-truth components by the fitted estimator under the three objectives, for the symmetric structure of (A). For each of 50 fits at $p = 1, 2, 4$, each recovered behavioral loading contributes one point, its absolute cosine similarity with the two true loadings, and the points lie near the unit quarter-circle. Separated solutions sit at the corners (open squares, the true components) and a fully mixed solution at $(1/\sqrt{2}, 1/\sqrt{2})$ (open diamond). The quartic objective ($p = 4$, red) recovers the true components, the linear objective ($p = 1$, blue) mixes them, and the squared objective ($p = 2$, gray) is flat in this symmetric case and scatters along the arc.

### Supplementary Note 2: Mathematical Formulation and Algorithm

#### Problem Formulation

##### Setup and Notation

For $K$ subjects with paired neural and behavioral recordings, each subject $k \in \{1, \ldots, K\}$ has neural data $\mathbf{X}_k \in \mathbb{R}^{T_k \times N_k}$ with $T_k$ time points and $N_k$ neurons, behavioral data $\mathbf{Y}_k \in \mathbb{R}^{T_k \times B}$ with $B$ behavioral features, a subject-specific neural projection $\mathbf{a}_k \in \mathbb{R}^{N_k}$, and a shared behavioral projection $\mathbf{b} \in \mathbb{R}^B$. We assume each column of $\mathbf{X}_k$ and $\mathbf{Y}_k$ has been mean-centered, so that the matrix products below give unbiased sample covariances.

The covariance matrices are defined using the unbiased estimator,

$$\mathbf{C}^k_{XX} = \frac{1}{T_k - 1}\mathbf{X}_k^\top \mathbf{X}_k \in \mathbb{R}^{N_k \times N_k} \quad \text{(neural covariance)}, \tag{16}$$

$$\mathbf{C}^k_{YY} = \frac{1}{T_k - 1}\mathbf{Y}_k^\top \mathbf{Y}_k \in \mathbb{R}^{B \times B} \quad \text{(behavioral covariance)}, \tag{17}$$

$$\mathbf{C}^k_{XY} = \frac{1}{T_k - 1}\mathbf{X}_k^\top \mathbf{Y}_k \in \mathbb{R}^{N_k \times B} \quad \text{(cross-covariance)}. \tag{18}$$

The average behavioral covariance across subjects is

$$\overline{\mathbf{C}_{YY}} = \frac{1}{K}\sum_{k=1}^{K} \mathbf{C}^k_{YY} \,. \tag{19}$$

##### Optimization Objective

ShaReD maximizes the sum of squared correlations between neural and behavioral projections across subjects, subject to unit variance constraints. The Lagrangian is

$$\mathcal{L} = -\sum_{k=1}^{K} \omega_k (\mathbf{a}_k^\top \mathbf{C}^k_{XY} \mathbf{b})^2 + \sum_{k=1}^{K} \alpha_k (\mathbf{a}_k^\top \mathbf{C}^k_{XX} \mathbf{a}_k - 1) + \beta(\mathbf{b}^\top \overline{\mathbf{C}_{YY}} \mathbf{b} - 1)\,, \tag{20}$$

where $\omega_k = 1/K$ weights each subject equally, and $\alpha_k$ and $\beta$ are Lagrange multipliers enforcing the unit variance constraints.

#### Derivation of the Solution

##### Optimizing Neural Projections

Setting the partial derivative with respect to $\mathbf{a}_k$ to zero,

$$\frac{\partial \mathcal{L}}{\partial \mathbf{a}_k} = -2\omega_k (\mathbf{a}_k^\top \mathbf{C}^k_{XY} \mathbf{b})(\mathbf{C}^k_{XY} \mathbf{b}) + 2\alpha_k \mathbf{C}^k_{XX} \mathbf{a}_k = 0\,, \tag{21}$$

and rearranging gives

$$\alpha_k \mathbf{C}^k_{XX} \mathbf{a}_k = \omega_k (\mathbf{a}_k^\top \mathbf{C}^k_{XY} \mathbf{b})(\mathbf{C}^k_{XY} \mathbf{b})\,. \tag{22}$$

Letting $O_k \equiv \mathbf{a}_k^\top \mathbf{C}^k_{XY} \mathbf{b}$ denote the overlap between projected neural and behavioral data for subject $k$, the formal solution for $\mathbf{a}_k$ is

$$\mathbf{a}_k = \frac{\omega_k O_k}{\alpha_k} (\mathbf{C}^k_{XX})^{-1} \mathbf{C}^k_{XY} \mathbf{b}\,. \tag{23}$$

**Optimizing Behavioral Projection**

Setting the partial derivative with respect to $\mathbf{b}$ to zero,

$$\frac{\partial \mathcal{L}}{\partial \mathbf{b}} = -2\sum_{k=1}^{K} \omega_k (\mathbf{a}_k^\top \mathbf{C}_{XY}^k \mathbf{b}) \mathbf{C}_{YX}^k \mathbf{a}_k + 2\beta \overline{\mathbf{C}_{YY}} \mathbf{b} = 0\,, \tag{24}$$

where $\mathbf{C}_{YX}^k = (\mathbf{C}_{XY}^k)^\top$. Substituting equation (23), we find

$$\sum_{k=1}^{K} \omega_k O_k \mathbf{C}_{YX}^k \left( \frac{\omega_k O_k}{\alpha_k} (\mathbf{C}_{XX}^k)^{-1} \mathbf{C}_{XY}^k \mathbf{b} \right) = \beta \overline{\mathbf{C}_{YY}} \mathbf{b}\,, \tag{25}$$

which simplifies to

$$\sum_{k=1}^{K} \frac{\omega_k^2 O_k^2}{\alpha_k} \mathbf{C}_{YX}^k (\mathbf{C}_{XX}^k)^{-1} \mathbf{C}_{XY}^k \mathbf{b} = \beta \overline{\mathbf{C}_{YY}} \mathbf{b}\,. \tag{26}$$

**Determining Lagrange Multipliers**

From the constraint $\mathbf{a}_k^\top \mathbf{C}_{XX}^k \mathbf{a}_k = 1$ and equation (23),

$$1 = \frac{\omega_k^2 O_k^2}{\alpha_k^2} \mathbf{b}^\top \mathbf{C}_{YX}^k (\mathbf{C}_{XX}^k)^{-1} \mathbf{C}_{XY}^k \mathbf{b}\,. \tag{27}$$

Substituting the expression for $\mathbf{a}_k$ into $O_k^2 = (\mathbf{a}_k^\top \mathbf{C}_{XY}^k \mathbf{b})^2$ and using the preceding equation gives

$$O_k^2 = \mathbf{b}^\top \mathbf{C}_{YX}^k (\mathbf{C}_{XX}^k)^{-1} \mathbf{C}_{XY}^k \mathbf{b}\,, \tag{28}$$

so that

$$\alpha_k = \omega_k O_k^2\,. \tag{29}$$

The constraint $\mathbf{b}^\top \overline{\mathbf{C}_{YY}} \mathbf{b} = 1$ combined with equation (26) gives

$$\beta = \sum_{k=1}^{K} \omega_k O_k^2\,. \tag{30}$$

**Final Eigenvalue Problem**

Substituting equation (29) into equation (26),

$$\sum_{k=1}^{K} \omega_k \mathbf{C}_{YX}^k (\mathbf{C}_{XX}^k)^{-1} \mathbf{C}_{XY}^k \mathbf{b} = \beta \overline{\mathbf{C}_{YY}} \mathbf{b}\,, \tag{31}$$

and multiplying both sides by $\overline{\mathbf{C}_{YY}}^{-1}$,

$$\sum_{k=1}^{K} \omega_k \overline{\mathbf{C}_{YY}}^{-1} \mathbf{C}_{YX}^k (\mathbf{C}_{XX}^k)^{-1} \mathbf{C}_{XY}^k \mathbf{b} = \beta \mathbf{b}\,. \tag{32}$$

The shared behavioral projection $\mathbf{b}$ is therefore an eigenvector of

$$\mathbf{M} = \sum_{k=1}^{K} \omega_k \overline{\mathbf{C}_{YY}}^{-1} \mathbf{C}_{YX}^k (\mathbf{C}_{XX}^k)^{-1} \mathbf{C}_{XY}^k\,, \tag{33}$$

with the leading component corresponding to the largest eigenvalue.

##### Incorporating Regularization

When behavioral features have temporal structure or when emphasizing consistently encoded features is desired, the objective is augmented with regularization terms,

$$\begin{aligned}\mathcal{L}_{reg} = &-\sum_{k=1}^{K}\omega_k(\mathbf{a}_k^\top \mathbf{C}_{XY}^k \mathbf{b})^2 + \sum_{k=1}^{K}\alpha_k(\mathbf{a}_k^\top \mathbf{C}_{XX}^k \mathbf{a}_k - 1) + \beta(\mathbf{b}^\top \overline{\mathbf{C}_{YY}}\mathbf{b} - 1) \\ &+ \lambda_{reg}\|\mathbf{b}\|_1 + \lambda_{smooth}\sum_{g=1}^{G}\sum_{j=2}^{n_g}(b_j^g - b_{j-1}^g)^2\,,\end{aligned} \tag{34}$$

where $\lambda_{reg}$ controls the L1 penalty on behavioral weights, $\lambda_{smooth}$ enforces smoothness within groups of temporally related features, $G$ is the number of behavioral variable groups, $n_g$ is the number of time lags for group $g$, and $b_j^g$ denotes the $j$-th weight in group $g$.

Because the L1 term is nonsmooth, ShaReD optimizes $\mathcal{L}_{reg}$ in two stages. Stage 1 solves the unregularized problem in closed form through the eigendecomposition of equation (33), which gives the global optimum of the squared-correlation objective and provides the initialization. Stage 2 refines this solution by proximal gradient descent, also called forward-backward splitting [27], taking gradient steps on the smooth part of the objective and applying the L1 penalty through a proximal soft-threshold. Let $\mathcal{L}_{diff} = \mathcal{L}_{reg} - \lambda_{reg}\|\mathbf{b}\|_1$ be that smooth part, consisting of every term except the L1 penalty. The smoothness penalty is differentiable and belongs to $\mathcal{L}_{diff}$.

##### Gradient Computations

The Stage 2 gradient steps act on $\mathcal{L}_{diff}$. Neither penalty depends on $\mathbf{a}_k$, so the gradient with respect to $\mathbf{a}_k$ is unchanged from the unregularized case,

$$\frac{\partial \mathcal{L}_{diff}}{\partial \mathbf{a}_k} = -2\omega_k(\mathbf{a}_k^\top \mathbf{C}_{XY}^k \mathbf{b})(\mathbf{C}_{XY}^k \mathbf{b}) + 2\alpha_k \mathbf{C}_{XX}^k \mathbf{a}_k\,. \tag{35}$$

The gradient with respect to $\mathbf{b}$ is

$$\frac{\partial \mathcal{L}_{diff}}{\partial \mathbf{b}} = -2\sum_{k=1}^{K}\omega_k(\mathbf{a}_k^\top \mathbf{C}_{XY}^k \mathbf{b})\mathbf{C}_{YX}^k \mathbf{a}_k + 2\beta\overline{\mathbf{C}_{YY}}\mathbf{b} + \nabla_{smooth}\,, \tag{36}$$

where $\nabla_{smooth}$ is the smoothness gradient derived below.

##### Smoothness Gradient with Boundary Conditions

The smoothness penalty $\sum_{g=1}^{G}\sum_{j=2}^{n_g}(b_j^g - b_{j-1}^g)^2$ requires careful treatment at group boundaries. The gradient depends on each element's position within its group.

**First element of group $g$ ($j = 1$),**

$$[\nabla_{smooth}]_1^g = 2\lambda_{smooth}(b_1^g - b_2^g)\,. \tag{37}$$

**Interior elements ($2 \le j \le n_g - 1$),**

$$[\nabla_{smooth}]_j^g = 2\lambda_{smooth}(2b_j^g - b_{j-1}^g - b_{j+1}^g)\,. \tag{38}$$

**Last element of group $g$ ($j = n_g$),**

$$[\nabla_{smooth}]_{n_g}^g = 2\lambda_{smooth}(b_{n_g}^g - b_{n_g-1}^g)\,. \tag{39}$$

**Single-element groups ($n_g = 1$),**

$$[\nabla_{smooth}]_1^g = 0\,. \tag{40}$$

The full smoothness gradient $\nabla_{smooth}$ is constructed by concatenating these group-wise gradients according to the feature ordering.

#### Two-Stage Optimization

**Stage 1, initialization via eigendecomposition.** Solve the unregularized problem (equation (33)) to obtain the initial $\mathbf{b}$, then compute the corresponding $\mathbf{a}_k$ values using the closed-form solution (equation (23)).

**Stage 2, refinement via gradient descent.** The variables satisfy the unit-variance constraints $\mathbf{a}_k^\top \mathbf{C}_{XX}^k \mathbf{a}_k = 1$ and $\mathbf{b}^\top \overline{\mathbf{C}_{YY}} \mathbf{b} = 1$, so each gradient step is taken in the ambient space and then retracted back onto the corresponding constraint surface. Each iteration $t$ updates $\mathbf{b}$ by a gradient step followed by the proximal soft-threshold,

$$\tilde{\mathbf{b}}^{(t+1)} = \mathbf{b}^{(t)} - \eta\, \partial \mathcal{L}_{diff} / \partial \mathbf{b}\,, \tag{41}$$

$$\mathbf{b}^{(t+1)} = \text{sign}(\tilde{\mathbf{b}}^{(t+1)}) \odot \max(|\tilde{\mathbf{b}}^{(t+1)}| - \eta \lambda_{reg}, 0)\,, \tag{42}$$

and retracts the result onto the behavioral constraint surface, $\mathbf{b} \leftarrow \mathbf{b}/\sqrt{\mathbf{b}^\top \overline{\mathbf{C}_{YY}} \mathbf{b}}$. The soft-threshold is the proximal operator of the L1 norm, $\text{prox}_{\eta\lambda_{reg}\|\cdot\|_1}(\tilde{\mathbf{b}}) = \arg\min_{\mathbf{b}} \left(\eta\lambda_{reg}\|\mathbf{b}\|_1 + \frac{1}{2}\|\mathbf{b} - \tilde{\mathbf{b}}\|_2^2\right)$. Each subject's neural projection is updated by gradient descent, $\mathbf{a}_k^{(t+1)} = \mathbf{a}_k^{(t)} - \eta\, \partial \mathcal{L}_{diff}/\partial \mathbf{a}_k$, and retracted onto its constraint surface, $\mathbf{a}_k \leftarrow \mathbf{a}_k / \sqrt{\mathbf{a}_k^\top \mathbf{C}_{XX}^k \mathbf{a}_k}$. The Lagrange multipliers are then set to their unit-variance values $\alpha_k = \omega_k (\mathbf{a}_k^\top \mathbf{C}_{XY}^k \mathbf{b})^2$ and $\beta = \sum_k \alpha_k$, and the learning rate decay schedule below is applied.

ShaReD extracts components sequentially by deflation (Multiple Component Extraction, below). Let $\hat{\mathbf{b}}^{(c)}$ and $\{\hat{\mathbf{a}}_k^{(c)}\}$ denote the unit-Euclidean-norm projections returned for component $c$, and $\{\hat{\mathbf{b}}^{(j)}\}_{j=1}^{c-1}$ the behavioral projections of the previously extracted components. Orthogonalization, deflation, and the returned vectors use these unit-Euclidean-norm versions ($\hat{\mathbf{b}}$, $\hat{\mathbf{a}}_k$) rather than the variance metric of the optimization. After the gradient descent loop completes, orthogonalization against previous components is applied once,

$$\mathbf{b}^{(c)} \leftarrow \hat{\mathbf{b}}^{(c)} - \sum_{j=1}^{c-1} (\hat{\mathbf{b}}^{(c)\top} \hat{\mathbf{b}}^{(j)}) \hat{\mathbf{b}}^{(j)}\,, \tag{43}$$

with analogous orthogonalizations for the neural projections. Each projection is then renormalized to unit Euclidean norm.

Algorithm 1 returns these unit-Euclidean-norm projections $\hat{\mathbf{a}}_k$ and $\hat{\mathbf{b}}$. The unit-variance projections of the derivation are recovered from them by rescaling in the covariance metric,

$$\mathbf{a}_k = \frac{\hat{\mathbf{a}}_k}{\sqrt{\hat{\mathbf{a}}_k^\top \mathbf{C}_{XX}^k \hat{\mathbf{a}}_k}}\,, \qquad \mathbf{b} = \frac{\hat{\mathbf{b}}}{\sqrt{\hat{\mathbf{b}}^\top \overline{\mathbf{C}_{YY}} \hat{\mathbf{b}}}}\,, \tag{44}$$

which restores the constraints $\mathbf{a}_k^\top \mathbf{C}_{XX}^k \mathbf{a}_k = 1$ and $\mathbf{b}^\top \overline{\mathbf{C}_{YY}} \mathbf{b} = 1$. The reported correlations and cosine similarities are unchanged by this choice of scale.

##### Learning Rate Schedule

Gradient descent runs for a fixed number of iterations. The learning rate decays exponentially,

$$\eta^{(t)} = \eta_0 \cdot \gamma^{\lfloor t/\tau \rfloor}, \tag{45}$$

where $\eta_0$ is the initial learning rate, $\gamma$ is the decay factor, and $\tau$ is the decay interval.

##### Complete Algorithm

**Algorithm 1** ShaReD Component Extraction

**Input:** Neural data $\{\mathbf{X}_k\}_{k=1}^K$, behavioral data $\{\mathbf{Y}_k\}_{k=1}^K$, regularization $\lambda_{reg}$, $\lambda_{smooth}$, component index $c$ with previously extracted projections $\{\hat{\mathbf{b}}^{(j)}, \hat{\mathbf{a}}_k^{(j)}\}_{j=1}^{c-1}$
**Output:** Behavioral projection $\hat{\mathbf{b}}$, neural projections $\{\hat{\mathbf{a}}_k\}_{k=1}^K$

// **Preprocessing.** $\mathbf{X}_k$, $\mathbf{Y}_k$ are z-scored and the behavioral matrix ZCA-whitened (see Implementation Details)

// **Compute covariances**
**for** $k = 1$ to $K$ **do**
  $\mathbf{C}_{XX}^k \leftarrow \frac{1}{T_k - 1}\mathbf{X}_k^\top \mathbf{X}_k + \epsilon \mathbf{I}, \quad \mathbf{C}_{YY}^k \leftarrow \frac{1}{T_k - 1}\mathbf{Y}_k^\top \mathbf{Y}_k, \quad \mathbf{C}_{XY}^k \leftarrow \frac{1}{T_k - 1}\mathbf{X}_k^\top \mathbf{Y}_k$
  $\omega_k \leftarrow 1/K$ // uniform
**end for**
$\overline{\mathbf{C}_{YY}} \leftarrow \frac{1}{K}\sum_{k=1}^K \mathbf{C}_{YY}^k + \epsilon \mathbf{I}$

// **Stage 1, eigendecomposition initialization**
$\mathbf{M} \leftarrow \sum_{k=1}^K \omega_k \overline{\mathbf{C}_{YY}}^{-1} \mathbf{C}_{YX}^k (\mathbf{C}_{XX}^k)^{-1} \mathbf{C}_{XY}^k$
$\mathbf{b} \leftarrow$ principal eigenvector of $\mathbf{M}$ (largest eigenvalue)
$\hat{\mathbf{b}} \leftarrow \mathbf{b}/\|\mathbf{b}\|_2$
**for** $k = 1$ to $K$ **do**
  $\mathbf{a}_k \leftarrow (\mathbf{C}_{XX}^k)^{-1}\mathbf{C}_{XY}^k \mathbf{b} \quad \hat{\mathbf{a}}_k \leftarrow \mathbf{a}_k/\|\mathbf{a}_k\|_2$
**end for**

// **Stage 2, gradient refinement (if** $\lambda_{reg} > 0$ **or** $\lambda_{smooth} > 0$**)**
**for** $t = 1$ to max_iterations **do**
  $\tilde{\mathbf{b}} \leftarrow \mathbf{b} - \eta \nabla_{\mathbf{b}} \mathcal{L}_{diff}$
  $\mathbf{b} \leftarrow \mathrm{sign}(\tilde{\mathbf{b}}) \odot \max(|\tilde{\mathbf{b}}| - \eta\lambda_{reg}, 0), \quad \mathbf{b} \leftarrow \mathbf{b}/\sqrt{\mathbf{b}^\top \overline{\mathbf{C}_{YY}} \mathbf{b}}$
  **for** $k = 1$ to $K$ **do**
    $\mathbf{a}_k \leftarrow \mathbf{a}_k - \eta \nabla_{\mathbf{a}_k} \mathcal{L}_{diff}, \quad \mathbf{a}_k \leftarrow \mathbf{a}_k/\sqrt{\mathbf{a}_k^\top \mathbf{C}_{XX}^k \mathbf{a}_k}$
    $\alpha_k \leftarrow \omega_k (\mathbf{a}_k^\top \mathbf{C}_{XY}^k \mathbf{b})^2$
  **end for**
  $\beta \leftarrow \sum_{k=1}^K \alpha_k$, apply learning rate decay
**end for**

$\mathbf{b} \leftarrow \hat{\mathbf{b}} - \sum_{j=1}^{c-1} (\hat{\mathbf{b}}^\top \hat{\mathbf{b}}^{(j)}) \hat{\mathbf{b}}^{(j)}$, similarly for each $\mathbf{a}_k$
$\hat{\mathbf{b}} \leftarrow \mathbf{b}/\|\mathbf{b}\|_2, \quad \hat{\mathbf{a}}_k \leftarrow \mathbf{a}_k/\|\mathbf{a}_k\|_2$

#### Multiple Component Extraction

Multiple components are extracted via deflation of the data.

**Algorithm 2** Multi-Component ShaReD with Deflation

```
1: for component c = 1 to n_components do
2:    Extract component c using Algorithm 1, store b̂^(c) and {â_k^(c)}
3:    for k = 1 to K do
4:       X_k ← X_k − (X_k â_k^(c))(â_k^(c))^⊤,    Y_k ← Y_k − (Y_k b̂^(c))(b̂^(c))^⊤
5:    end for
6: end for
```

Bilateral deflation removes variance along previous components from both $\mathbf{X}_k$ and $\mathbf{Y}_k$, so the Stage 1 eigendecomposition for component $c$ produces projections that are orthogonal to all previous components. The explicit Gram-Schmidt orthogonalization step in Algorithm 1 is needed only to restore this orthogonality after Stage 2 gradient refinement, and has no effect when refinement is not used.

#### Determining the Number of Components

Each extracted component is tested for significance by cross-validation against a permutation null. ShaReD is fit on training data, test-set correlations are computed per component, and each is compared against a null obtained by refitting on shuffled data. The significant count is the size of the leading contiguous sequence of components that pass significance testing with Benjamini-Hochberg correction ($q < 0.05$), and the number of components carried into a downstream analysis is chosen at or below it.

#### Numerical Considerations

##### Matrix Regularization

Covariance matrices may be ill-conditioned, particularly with limited data or highly correlated features. A small regularization term is added to the diagonal of $\mathbf{C}_{XX}^k$ and $\overline{\mathbf{C}_{YY}}$, $\mathbf{C}_{XX}^k \leftarrow \mathbf{C}_{XX}^k + \epsilon\mathbf{I}$ and $\overline{\mathbf{C}_{YY}} \leftarrow \overline{\mathbf{C}_{YY}} + \epsilon\mathbf{I}$.

##### Handling Degenerate Solutions

In rare cases, the eigendecomposition may yield degenerate solutions, for example when behavioral features are nearly linearly dependent. Degeneracy is detected when the top eigenvalues are nearly equal, in which case the leading direction is a rotational mixture, or when the resulting correlations are near zero. The latter is equivalent to the per-subject neural-behavioral overlaps $O_k = \mathbf{a}_k^\top \mathbf{C}_{XY}^k \mathbf{b}$ being near zero for every subject, since the leading eigenvalue equals $\sum_k \omega_k O_k^2$ (equation (30)). A component with no consistent neural correlate in any subject carries no shared signal, so its direction is set by noise rather than by the data and is not identifiable. When detected, increasing regularization or reducing the behavioral feature dimensionality typically resolves the issue.

##### Computational Complexity

The per-component complexity comprises covariance computation $\mathcal{O}(\sum_k T_k(N_k^2 + B^2))$, matrix inversions $\mathcal{O}(\sum_k N_k^3)$, calculation of the contributions to $\mathbf{M}$ of each subject $\mathcal{O}(\sum_k N_k B^2)$, eigendecomposition of $\mathbf{M}$ with complexity $\mathcal{O}(B^3)$, per-iteration gradient descent $\mathcal{O}(\sum_k N_k B)$, and deflation $\mathcal{O}(\sum_k T_k(N_k + B))$.

The total complexity for $C$ components with $I$ gradient iterations is

$$\mathcal{O}\left(C \cdot \left[\sum_k T_k(N_k^2 + B^2) + \sum_k N_k^3 + \sum_k N_k B^2 + B^3 + I \sum_k N_k B\right]\right). \tag{46}$$

#### Supplementary Note 3: Synthetic Data Generation and Validation

This note describes the synthetic data generation procedure used to validate ShaReD, including the parameterization that controls neural-behavioral correlations, the sample-size sensitivity analysis, and the different sharing patterns across subjects.

##### Generative Model

###### Basic Model

Paired neural and behavioral observations are generated from shared latent factors. For $K$ subjects, each with $C$ components, the generative model is

$$\mathbf{X}_k = \mathbf{Z}\boldsymbol{\alpha}\mathbf{A}_k^\top + \boldsymbol{\varepsilon}_X^k \,, \tag{47}$$

$$\mathbf{Y}_k = \mathbf{Z}\boldsymbol{\beta}\mathbf{B}^\top + \boldsymbol{\varepsilon}_Y^k \,, \tag{48}$$

where $\mathbf{Z} \in \mathbb{R}^{T\times C}$ contains $C$ latent factors over $T$ time points, with columns drawn i.i.d. from $\mathcal{N}(0, \mathbf{I}_T)$ and then SVD-orthonormalized to $\|\mathbf{Z}_{:,c}\| = \sqrt{T}$, so that distinct components are exactly orthogonal. $\mathbf{A}_k \in \mathbb{R}^{N_k\times C}$ contains subject-specific neural loadings, $\mathbf{B} \in \mathbb{R}^{B\times C}$ contains shared behavioral loadings, $\boldsymbol{\alpha}, \boldsymbol{\beta} \in \mathbb{R}^{C\times C}$ are diagonal scaling matrices controlling signal strength, and $\boldsymbol{\varepsilon}_X^k \sim \mathcal{N}(0, \sigma^2\mathbf{I})$ and $\boldsymbol{\varepsilon}_Y^k \sim \mathcal{N}(0, \sigma^2\mathbf{I})$ add independent noise.

###### Generating Orthonormal Loadings

Ground-truth loading matrices are generated by drawing random matrices $\tilde{\mathbf{A}}_k \in \mathbb{R}^{N_k\times C}$ and $\tilde{\mathbf{B}} \in \mathbb{R}^{B\times C}$ with i.i.d. $\mathcal{N}(0, 1)$ entries, then orthonormalizing via QR decomposition, $\mathbf{A}_k = \mathrm{orth}(\tilde{\mathbf{A}}_k)$ and $\mathbf{B} = \mathrm{orth}(\tilde{\mathbf{B}})$. This ensures that different components are orthogonal and that the scaling matrices $\boldsymbol{\alpha}$ and $\boldsymbol{\beta}$ directly control signal magnitude.

##### Controlling Neural-Behavioral Correlations

###### Target Correlation Specification

For each component $c$, a target correlation $\rho_c^*$ is specified between the projected neural and behavioral data. The scaling factors $\alpha_c$ and $\beta_c$ are determined such that

$$\mathrm{Corr}(\mathbf{X}_k\mathbf{a}_k^{(c)}, \mathbf{Y}_k\mathbf{b}^{(c)}) = \rho_c^* \,, \tag{49}$$

where $\mathbf{a}_k^{(c)}$ and $\mathbf{b}^{(c)}$ are the true loading vectors for component $c$.

###### Derivation of Scaling Factors

For component $c$, the projected data are

$$\mathbf{x}_c^k = \mathbf{X}_k\mathbf{a}_k^{(c)} = \alpha_c\mathbf{z}_c + \boldsymbol{\varepsilon}_X^k\mathbf{a}_k^{(c)} \,, \tag{50}$$

$$\mathbf{y}_c = \mathbf{Y}_k\mathbf{b}^{(c)} = \beta_c\mathbf{z}_c + \boldsymbol{\varepsilon}_Y^k\mathbf{b}^{(c)} \,, \tag{51}$$

where $\mathbf{z}_c = \mathbf{Z}_{:,c}$ is the $c$-th latent factor.

The covariance between projected data is

$$\mathrm{Cov}(\mathbf{x}_c^k, \mathbf{y}_c) = \alpha_c\beta_c\mathrm{Var}(\mathbf{z}_c) = \alpha_c\beta_c \,, \tag{52}$$

and the variances are

$$\mathrm{Var}(\mathbf{x}_c^k) = \alpha_c^2 + \sigma^2 \|\mathbf{a}_k^{(c)}\|^2 = \alpha_c^2 + \sigma^2 \,, \tag{53}$$

$$\mathrm{Var}(\mathbf{y}_c) = \beta_c^2 + \sigma^2 \|\mathbf{b}^{(c)}\|^2 = \beta_c^2 + \sigma^2 \,. \tag{54}$$

The resulting correlation is

$$\rho_c = \frac{\alpha_c \beta_c}{\sqrt{(\alpha_c^2 + \sigma^2)(\beta_c^2 + \sigma^2)}} \,. \tag{55}$$

##### Symmetric Scaling Solution

Using symmetric scaling $\alpha_c = \beta_c = s_c$, the correlation becomes

$$\rho_c = \frac{s_c^2}{s_c^2 + \sigma^2} \,. \tag{56}$$

Solving for $s_c$ given target correlation $\rho_c^*$,

$$s_c = \sigma \sqrt{\frac{\rho_c^*}{1 - \rho_c^*}} \,. \tag{57}$$

This formula is valid for $0 < \rho_c^* < 1$. For $\rho_c^* = 0$, set $s_c = 0$ (pure noise). As $\rho_c^* \to 1$, $s_c \to \infty$, since perfect correlation requires infinite signal-to-noise ratio.

##### Practical Implementation

Given target correlations $\{\rho_1^*, \rho_2^*, \dots, \rho_C^*\}$ and noise level $\sigma$, scaling factors are computed as $s_c = \sigma\sqrt{\rho_c^*/(1 - \rho_c^*)}$ for each $c$, the diagonal matrices are constructed as $\boldsymbol{\alpha} = \boldsymbol{\beta} = \mathrm{diag}(s_1, s_2, \dots, s_C)$, data are generated using the above generative model, and the achieved correlations are verified against targets up to finite-sample variation.

#### Preprocessing

Synthetic data were preprocessed in the same way as the neural and behavioral recordings before ShaReD was applied. Each feature was z-scored to zero mean and unit variance, and then both $\mathbf{X}$ and $\mathbf{Y}$ were whitened. The signal-strength, pseudopopulation, identifiability, and extended sample-size sweeps used ZCA whitening, with $\mathbf{W} = \mathbf{V}\,\mathrm{diag}\big(\max(\lambda_i, \epsilon)\big)^{-1/2}\,\mathbf{V}^\top$ and eigenvalue floor $\epsilon = 10^{-8}$, obtained from the eigendecomposition of the training-set sample covariance and applied separately to training and test data to avoid leakage. The sharing-structure analysis whitened on the full dataset using the same ZCA transform, computed in SVD form. Because recovery is scored against the ground-truth loadings rather than a held-out correlation, full-dataset whitening introduces no leakage.

#### Component Recovery Assessment

##### Matching Recovered to True Components

ShaReD may recover components in a different order than the ground truth. Recovered components were matched to true components using the Hungarian algorithm based on cosine similarity,

$$\mathrm{sim}_{ij} = \frac{|\check{\mathbf{b}}^{(i)\top}\mathbf{b}^{(j)}|}{\|\check{\mathbf{b}}^{(i)}\|\|\mathbf{b}^{(j)}\|} \,, \tag{58}$$

where $\check{\mathbf{b}}^{(i)}$ is the $i$-th recovered behavioral projection and $\mathbf{b}^{(j)}$ is the $j$-th true behavioral projection. The absolute value accounts for sign ambiguity, since projections are equivalent up to sign flip.

**Recovery Metrics**

For each matched pair $(i, j)$, behavioral projection recovery is

$$\text{recovery}_b^{(j)} = \frac{|\check{\mathbf{b}}^{(i)\top}\mathbf{b}^{(j)}|}{\|\check{\mathbf{b}}^{(i)}\|\|\mathbf{b}^{(j)}\|}, \tag{59}$$

neural projection recovery, averaged across subjects, is

$$\text{recovery}_a^{(j)} = \frac{1}{K}\sum_{k=1}^{K} \frac{|\check{\mathbf{a}}_k^{(i)\top}\mathbf{a}_k^{(j)}|}{\|\check{\mathbf{a}}_k^{(i)}\|\|\mathbf{a}_k^{(j)}\|}, \tag{60}$$

and the test correlation is

$$\rho_{test}^{(j)} = \frac{1}{K}\sum_{k=1}^{K} \left|\text{Corr}(\mathbf{X}_k^{test}\check{\mathbf{a}}_k^{(i)}, \mathbf{Y}_k^{test}\check{\mathbf{b}}^{(i)})\right|. \tag{61}$$

#### Standard Validation Configuration

Unless otherwise noted, synthetic experiments used the parameters in Table S1.

| **Parameter** | **Value** |
|---|---|
| Number of subjects ($K$) | 6 |
| Neural features per subject ($N_k$) | 50 |
| Behavioral features ($B$) | 50 |
| Training samples ($T_{train}$) | 20,000 |
| Test samples ($T_{test}$) | 5,000 |
| Noise standard deviation ($\sigma$) | 1.0 |
| Number of components ($C$) | 14 |
| Target correlations | 0.70, 0.65, . . . , 0.10, 0.05 |
| Random seeds | 100 repetitions |

Table S1: Default synthetic data parameters for signal-strength analysis.

#### Identifiability of Nearby Components versus Single-Subject CCA

Each dataset contained two components in paired $\mathbf{X}$ and $\mathbf{Y}$ views with 50 features per view. One component was anchored at canonical correlation $\rho_1$, and the second sat at $\rho_1 + \Delta$ with signed offset $\Delta$. Gaussian noise with $\sigma = 1$ was added to both views. ShaReD was fit with $K = 6$ subjects, $p = 2$, and 100 iterations to extract two components. Single-subject CCA was fit with $K = 1$ and two components. Recovered behavioral loadings were matched to ground truth by Hungarian assignment, and the cosine similarity between the recovered loading and the true loading of the anchor component was averaged over 100 repetitions.

Two CCA baselines were used. Under matched per-subject $T$, the CCA fit saw the same sample count as one ShaReD subject, so $T_{total}^{CCA} = T_{per-subj}^{ShaReD}$. Under matched total $T$, the CCA fit saw the combined sample count of all ShaReD subjects in a single subject, so $T_{total}^{CCA} = K \cdot T_{per-subj}^{ShaReD}$. Two complementary sweeps were run for each baseline. The first fixed $T_{total}^{ShaReD} = 12{,}000$ and varied $\rho_1 \in \{0.1, 0.2, \ldots, 0.8\}$. The second fixed $\rho_1 = 0.5$ and varied $T_{total}^{ShaReD} \in$ {1,200, 3,000, 6,000, 12,000, 30,000, 60,000}. Identifiability was assessed by absolute cosine similarity between the recovered and true behavioral loading of the anchor component, with Hungarian matching of recovered to true components [10].

#### Objective Function Comparison

The end-to-end recovery test (Fig. S1D) used the two-subject symmetric structure of Supplementary Note 1, with per-subject correlations $(0.9, 0.5)$ and $(0.5, 0.9)$, equal weights $\omega_k = 1/2$, 50 neural and behavioral features per subject, $T = 3{,}000$ samples, and noise $\sigma = 1.0$. Data were generated, z-scored, and whitened on the full dataset by the SVD form of the ZCA transform, as for the sharing-structure analysis, since the recovery test has no held-out fold. For each of 50 random-seed repetitions, ShaReD was fit at $p = 1, 2, 4$. All three fits were initialized from the $p = 2$ eigenvector and refined by 8,000 gradient steps on their own objective, with learning rate 0.05 and no regularization. Recovery was scored as the absolute cosine similarity of each recovered behavioral loading with each of the two true loadings, giving the two coordinates of each point on the quarter-circle.

#### Sample Size Sensitivity Analysis

To characterize how ShaReD performance scales with data quantity, we evaluated component recovery and test correlations across training set sizes.

##### Experimental Design

Training set sizes were swept over $T_{train} \in \{100, 200, 500, 1{,}000, 2{,}000, 5{,}000, 10{,}000\}$ with all other parameters at their Table S1 defaults.

##### Results Summary

Recovery quality depended on both sample size and target correlation strength. Strong components ($\rho^* \geq 0.5$) were reliably recovered (cosine similarity $> 0.9$) from $T \geq 500$ samples, moderate components ($0.2 \leq \rho^* < 0.5$) required $T \geq 2{,}000$ samples, and the weakest component ($\rho^* = 0.05$) was not reliably recovered at any tested sample size.

#### Components with Different Sharing Structure

To test ShaReD's ability to discover components with varying degrees of sharing, we generated data in which different components are shared among different subsets of subjects.

##### Sharing-Structure Generative Model

The basic model extends to allow component-specific subject subsets,

$$\mathbf{X}_k = \sum_{c=1}^{C} \mathbf{1}[k \in \mathcal{S}_c] \cdot \mathbf{z}_c \alpha_{c,k} \mathbf{a}_k^{(c)\top} + \boldsymbol{\varepsilon}_X^k \,, \tag{62}$$

$$\mathbf{Y}_k = \sum_{c=1}^{C} \mathbf{1}[k \in \mathcal{S}_c] \cdot \mathbf{z}_c \beta_c \mathbf{b}^{(c)\top} + \boldsymbol{\varepsilon}_Y^k \,, \tag{63}$$

where $\mathcal{S}_c \subseteq \{1, \ldots, K\}$ is the set of subjects that share component $c$, $\mathbf{1}[\cdot]$ is the indicator function, $\alpha_{c,k}$ is the per-subject neural-side scaling, and $\beta_c$ is the per-component behavioral-side scaling. The scaling factors are specified in the Generation Procedure below.

##### Five-Component Example

Seven subjects were generated with five components having distinct sharing patterns (Table S2).

| **Component** | **Subjects** | **Support** |
|---|---|---|
| $G$ (Global) | 1, 2, 3, 4, 5, 6, 7 | all subjects |
| $S_A$ (Subgroup A) | 1, 2, 3, 4 | contiguous block |
| $S_B$ (Subgroup B) | 5, 6, 7 | contiguous block |
| $S_C$ (Pair) | 4, 5 | bridges $S_A$ and $S_B$ |
| $S_D$ (Pair) | 2, 3 | lies within $S_A$ |

Table S2: Component sharing structure. All components target $\bar{\rho} = 0.60$, with per-subject correlations $\rho_{c,k}$ drawn independently from $\mathcal{N}(0.60, 0.03^2)$, clipped to $(0.05, 0.95)$, and resampled on each repetition. The two pair components have identical subject counts but distinct support, with $S_C$ bridging the two subgroups and $S_D$ lying entirely within $S_A$.

##### Sharing-Structure Validation Parameters

The sharing-structure experiments used $K = 7$ subjects, $N_k = B = 50$ neural and behavioral features, $T = 30{,}000$ training samples, noise level $\sigma = 1.0$, 1,000 optimization iterations, and 100 random-seed repetitions.

##### Generation Procedure

For each component $c$, the sharing set $\mathcal{S}_c$ was defined and component-specific latent factors $\mathbf{z}_c$ were generated. Per-subject target correlations $\rho_{c,k}$ were drawn from $\mathcal{N}(0.60, 0.03^2)$ clipped to $(0.05, 0.95)$ for $k \in \mathcal{S}_c$. Neural-side scaling used the per-subject target, $\alpha_{c,k} = \sigma\sqrt{\rho_{c,k}/(1-\rho_{c,k})}$, while behavioral-side scaling used the per-component mean $\bar{\rho}_c = (1/|\mathcal{S}_c|)\sum_{k\in\mathcal{S}_c} \rho_{c,k}$ as $\beta_c = \sigma\sqrt{\bar{\rho}_c/(1-\bar{\rho}_c)}$. The achieved per-subject correlation was $\sqrt{\rho_{c,k}\,\bar{\rho}_c}$, which reduces to $\rho_{c,k}$ when subjects in $\mathcal{S}_c$ share the same target and stays within a few percent of $\rho_{c,k}$ (median ∼1.4%) for the per-subject variability used here. Neural loadings $\mathbf{a}_k^{(c)}$ were drawn for subjects in $\mathcal{S}_c$ and set to zero for subjects outside it. Shared behavioral loadings $\mathbf{b}^{(c)}$ were generated for each component, and the data matrices were constructed by summing contributions from all components.

##### Validation Approach

ShaReD was applied without any information about the sharing structure. Recovery was assessed by component identification, by the cosine similarity between recovered and true behavioral projections, by the off-diagonal elements of the similarity matrix, and by whether recovered neural weights showed the zero/nonzero structure matching the sharing sets.

This recovery depends on the squared correlation objective. A linear objective can mix components with similar aggregate correlation strengths across subjects, while the squared objective never prefers a mixture (Supplementary Note 1).

#### Pseudopopulation Decoding Validation

Pseudopopulations were used to test whether recovered components provide a common reference frame for pooling neural data across subjects (Fig. 1D). Because the data were generated from ShaReD's own linear model, this analysis is a recovery test rather than an independent validation.

##### Parameters

The decoding analysis used 10,000 training samples per subject, 2,000 test samples per subject, 2,000 pseudopopulation time points, and a logistic regression classifier with up to 1,000 iterations, across 100 random-seed repetitions.

##### Procedure

Pseudopopulations were constructed by concatenating neural activity $[\mathbf{X}_1(t,:), \mathbf{X}_2(t,:), \ldots, \mathbf{X}_K(t,:)]$ across subjects at each time point, which simulated a large simultaneous recording under independent subject noise. Behavioral labels were defined by projecting behavioral data onto each recovered component direction, $\mathbf{y}_c(t) = \mathbf{Y}_k(t,:)\check{\mathbf{b}}^{(c)}$, and binarizing by median split. A logistic regression classifier was trained on pseudopopulation neural data to predict the binary label, and evaluated on a held-out pseudopopulation. Baselines included shuffled labels (chance), random projection directions in behavioral space, and individual raw behavioral feature dimensions.

#### Reproducibility

Synthetic experiments used base random seed 100, with per-repetition seeds computed as seed + repetition_index, except the parameter-sweep experiments, which offset the seed by grid cell so that different sweep conditions draw independent data.

#### Supplemental Figures

Table S3: **Per-session recording metadata and inclusion status.** Each row corresponds to one recording session from one task (CO or RT). "Neurons" is the unit count after preprocessing (firing-rate filtering at ≥1 Hz, with all units assigned to M1). "Min trials/target" is the minimum trial count across the 8 center-out targets (applicable to CO only, since RT uses random, non-discrete targets). Sessions were included if they had ≥30 neurons and, for CO, ≥20 trials per target. Subject J (3 CO sessions, 18–38 neurons) was excluded entirely. Two additional sessions fell below the trial-count threshold with unbalanced designs: M-2014-06-26 (6 trials for one target out of 1,198 total) and T-2013-08-19 (18 trials for one target). M-2014-06-27 fell below the neuron threshold (24 neurons after preprocessing). T-2013-08-30 fell below the neuron threshold (27 neurons).

| Session ID | Subject | Task | Neurons | Trials | Min trials/tgt | Status |
|---|---|---|---|---|---|---|
| C-2013-10-03 | C | CO | 57 | 193 | 19 | Excluded[a] |
| C-2013-10-09 | C | RT | 71 | 187 | – | Included |
| C-2013-10-10 | C | RT | 38 | 193 | – | Included |
| C-2013-10-11 | C | RT | 77 | 175 | – | Included |
| C-2013-10-22 | C | CO | 34 | 182 | 17 | Excluded[a] |
| C-2013-10-23 | C | CO | 47 | 224 | 21 | Included |
| C-2013-10-28 | C | RT | 55 | 174 | – | Included |
| C-2013-10-29 | C | RT | 47 | 194 | – | Included |
| C-2013-10-31 | C | CO | 42 | 247 | 26 | Included |
| C-2013-11-01 | C | CO | 33 | 280 | 30 | Included |
| C-2013-12-03 | C | CO | 46 | 208 | 17 | Excluded[a] |
| C-2013-12-04 | C | CO | 38 | 178 | 20 | Included |
| C-2013-12-09 | C | RT | 50 | 205 | – | Included |
| C-2013-12-10 | C | RT | 41 | 203 | – | Included |
| C-2013-12-12 | C | RT | 34 | 198 | – | Included |
| C-2013-12-13 | C | RT | 47 | 198 | – | Included |
| C-2013-12-17 | C | RT | 47 | 197 | – | Included |
| C-2013-12-18 | C | RT | 49 | 188 | – | Included |
| C-2013-12-19 | C | CO | 52 | 212 | 21 | Included |
| C-2013-12-20 | C | CO | 52 | 233 | 24 | Included |
| C-2015-03-09 | C | CO | 72 | 788 | 77 | Included |
| C-2015-03-11 | C | CO | 86 | 1,067 | 118 | Included |
| C-2015-03-12 | C | CO | 90 | 1,058 | 115 | Included |
| C-2015-03-13 | C | CO | 85 | 1,179 | 131 | Included |
| C-2015-03-16 | C | RT | 74 | 654 | – | Included |
| C-2015-03-17 | C | RT | 77 | 694 | – | Included |
| C-2015-03-18 | C | RT | 74 | 921 | – | Included |
| C-2015-03-19 | C | CO | 71 | 1,155 | 130 | Included |
| C-2015-03-20 | C | RT | 66 | 841 | – | Included |
| C-2015-06-29 | C | CO | 48 | 295 | 29 | Included |
| C-2015-06-30 | C | CO | 43 | 243 | 26 | Included |
| C-2015-07-01 | C | CO | 48 | 282 | 30 | Included |

*Continued on next page*

*Table S3 continued*

| Session ID | Subject | Task | Neurons | Trials | Min trials/tgt | Status |
|---|---|---|---|---|---|---|
| C-2015-07-03 | C | CO | 50 | 229 | 24 | Included |
| C-2015-07-06 | C | CO | 45 | 221 | 24 | Included |
| C-2015-07-07 | C | CO | 41 | 222 | 24 | Included |
| C-2015-07-08 | C | CO | 63 | 214 | 24 | Included |
| C-2015-07-09 | C | CO | 60 | 218 | 25 | Included |
| C-2015-07-10 | C | CO | 53 | 231 | 22 | Included |
| C-2015-07-13 | C | CO | 56 | 218 | 23 | Included |
| C-2015-07-14 | C | CO | 64 | 227 | 25 | Included |
| C-2015-07-15 | C | CO | 57 | 212 | 24 | Included |
| C-2015-07-16 | C | CO | 59 | 207 | 24 | Included |
| C-2015-11-03 | C | CO | 37 | 320 | 33 | Included |
| C-2015-11-04 | C | CO | 58 | 440 | 45 | Included |
| C-2015-11-06 | C | CO | 59 | 303 | 32 | Included |
| C-2015-11-09 | C | CO | 64 | 210 | 21 | Included |
| C-2015-11-10 | C | CO | 60 | 219 | 23 | Included |
| C-2015-11-12 | C | CO | 37 | 202 | 22 | Included |
| C-2015-11-13 | C | CO | 52 | 200 | 22 | Included |
| C-2015-11-16 | C | CO | 52 | 201 | 21 | Included |
| C-2015-11-17 | C | CO | 57 | 221 | 23 | Included |
| C-2015-11-19 | C | CO | 59 | 226 | 26 | Included |
| C-2015-11-20 | C | CO | 54 | 198 | 22 | Included |
| C-2015-12-01 | C | CO | 45 | 219 | 24 | Included |
| C-2016-09-09 | C | CO | 216 | 335 | 23 | Included |
| C-2016-09-12 | C | CO | 229 | 326 | 23 | Included |
| C-2016-09-14 | C | CO | 337 | 322 | 18 | Excluded[a] |
| C-2016-09-15 | C | CO | 295 | 291 | 22 | Included |
| C-2016-09-19 | C | CO | 281 | 276 | 23 | Included |
| C-2016-09-21 | C | CO | 283 | 271 | 24 | Included |
| C-2016-09-23 | C | CO | 276 | 278 | 16 | Excluded[a] |
| C-2016-09-29 | C | CO | 179 | 253 | 26 | Included |
| C-2016-10-05 | C | CO | 232 | 252 | 25 | Included |
| C-2016-10-06 | C | CO | 240 | 251 | 25 | Included |
| C-2016-10-07 | C | CO | 198 | 214 | 22 | Included |
| C-2016-10-11 | C | CO | 224 | 239 | 21 | Included |
| C-2016-10-13 | C | CO | 206 | 271 | 30 | Included |
| C-2016-10-21 | C | CO | 279 | 331 | 35 | Included |
| J-2016-04-05 | J | CO | 38 | 322 | – | Excluded[b] |
| J-2016-04-06 | J | CO | 18 | 296 | – | Excluded[b] |
| J-2016-04-07 | J | CO | 19 | 331 | – | Excluded[b] |
| M-2014-01-14 | M | RT | 133 | 252 | – | Included |
| M-2014-01-15 | M | RT | 139 | 246 | – | Included |
| M-2014-01-16 | M | RT | 149 | 198 | – | Included |
| M-2014-02-03 | M | CO | 78 | 245 | 23 | Included |

*Continued on next page*

*Table S3 continued*

| Session ID | Subject | Task | Neurons | Trials | Min trials/tgt | Status |
|---|---|---|---|---|---|---|
| M-2014-02-14 | M | RT | 96 | 201 | – | Included |
| M-2014-02-17 | M | CO | 114 | 255 | 25 | Included |
| M-2014-02-18 | M | CO | 124 | 259 | 28 | Included |
| M-2014-02-21 | M | RT | 109 | 209 | – | Included |
| M-2014-02-24 | M | RT | 100 | 182 | – | Included |
| M-2014-03-03 | M | CO | 96 | 233 | 22 | Included |
| M-2014-03-04 | M | CO | 105 | 235 | 23 | Included |
| M-2014-03-06 | M | CO | 116 | 264 | 25 | Included |
| M-2014-03-07 | M | CO | 85 | 254 | 28 | Included |
| M-2014-06-26 | M | CO | 25 | 1,198 | 6 | Excluded[a,c] |
| M-2014-06-27 | M | CO | 24 | 1,250 | 0 | Excluded[c] |
| M-2014-09-29 | M | CO | 112 | 805 | 94 | Included |
| M-2014-12-03 | M | CO | 55 | 223 | 25 | Included |
| M-2015-05-11 | M | CO | 64 | 621 | 66 | Included |
| M-2015-05-12 | M | CO | 84 | 563 | 55 | Included |
| M-2015-06-10 | M | CO | 47 | 208 | 24 | Included |
| M-2015-06-11 | M | CO | 46 | 215 | 23 | Included |
| M-2015-06-12 | M | CO | 36 | 206 | 22 | Included |
| M-2015-06-15 | M | CO | 38 | 205 | 20 | Included |
| M-2015-06-16 | M | CO | 50 | 211 | 22 | Included |
| M-2015-06-17 | M | CO | 49 | 223 | 20 | Included |
| M-2015-06-23 | M | CO | 49 | 207 | 21 | Included |
| M-2015-06-25 | M | CO | 57 | 219 | 21 | Included |
| M-2015-06-26 | M | CO | 53 | 208 | 21 | Included |
| T-2013-08-19 | T | CO | 48 | 179 | 18 | Excluded[a] |
| T-2013-08-20 | T | RT | 33 | 154 | – | Included |
| T-2013-08-21 | T | CO | 52 | 197 | 20 | Included |
| T-2013-08-22 | T | RT | 52 | 247 | – | Included |
| T-2013-08-23 | T | CO | 42 | 256 | 25 | Included |
| T-2013-08-30 | T | RT | 27 | 204 | – | Excluded[c] |
| T-2013-09-03 | T | CO | 45 | 216 | 22 | Included |
| T-2013-09-04 | T | RT | 65 | 162 | – | Included |
| T-2013-09-05 | T | CO | 58 | 212 | 22 | Included |
| T-2013-09-06 | T | RT | 42 | 216 | – | Included |
| T-2013-09-09 | T | CO | 50 | 280 | 26 | Included |
| T-2013-09-10 | T | RT | 59 | 229 | – | Included |

[a]Min trials per target below 20 threshold.

[b]Subject excluded entirely (all sessions 18–38 neurons).

[c]Neurons below 30 threshold.

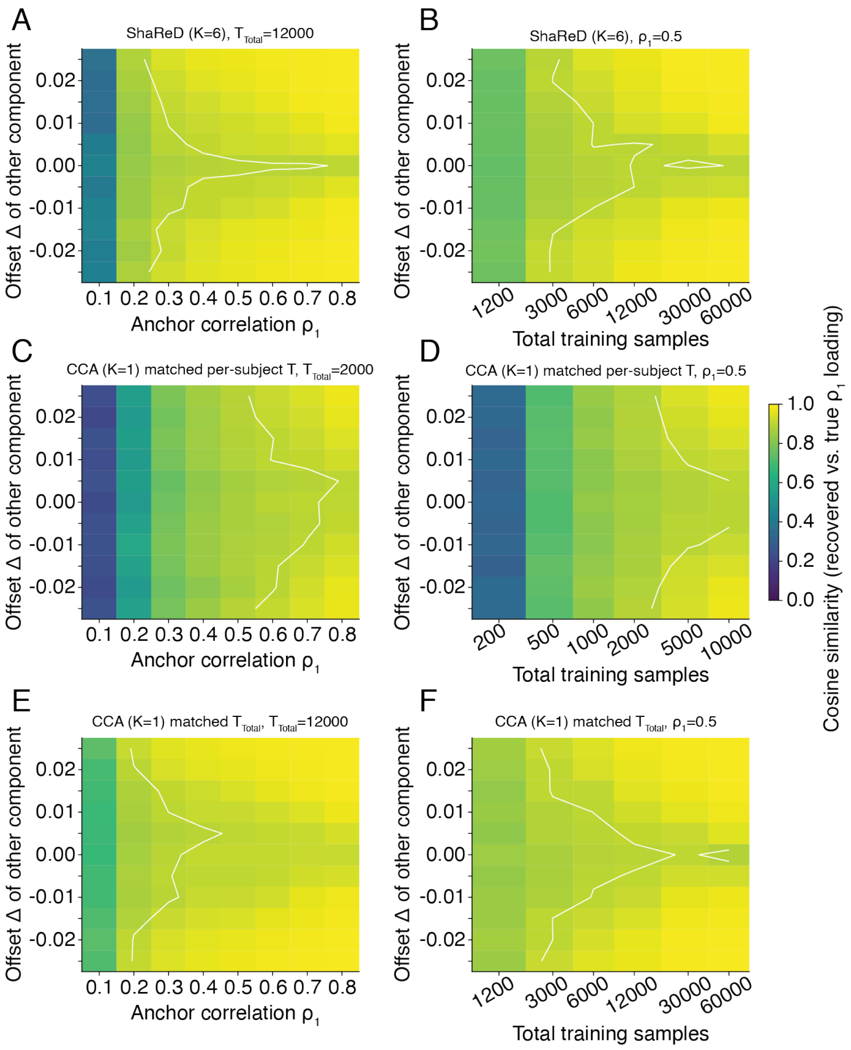


Figure S2: **Identifiability of nearby components against single-subject CCA.** ShaReD ($K = 6$) is compared to single-subject CCA ($K = 1$) under two matching schemes. Each dataset contains two components, one anchored at canonical correlation $\rho_1$ and the other at $\rho_1 + \Delta$. The $y$-axis shows the signed offset $\Delta$ and color shows the cosine similarity between the recovered loading and the true loading of the anchor component (mean across 100 repetitions, with Hungarian matching of recovered to true components). White contours mark cosine similarity $= 0.9$. **(A)** ShaReD recovery as a function of anchor correlation $\rho_1$ at $T_{total} = 12{,}000$ (per-subject $T = 2{,}000$). **(B)** ShaReD recovery as a function of total training samples at $\rho_1 = 0.5$. **(C, D)** CCA at matched per-subject $T$, where the CCA fit sees the same sample count as one ShaReD subject. (C) varies $\rho_1$ at $T_{total} = 2{,}000$. (D) varies $T_{total}$ at $\rho_1 = 0.5$. **(E, F)** CCA at matched total $T$, where the CCA fit sees the same total sample count as all ShaReD subjects combined, in a single subject. (E) varies $\rho_1$ at $T_{total} = 12{,}000$. (F) varies $T_{total}$ at $\rho_1 = 0.5$. Pooling several lightly sampled subjects outperforms CCA on any single one of them (A versus C, B versus D). With the total sample count held fixed, ShaReD recovers nearly as much from many subjects as CCA does from one subject providing all the samples (A versus E, B versus F), and trails only near the tie at $\Delta = 0$, where equal correlation values leave the anchor component unidentifiable for any method. This degeneracy is inherited from CCA and appears as the narrow spike at small $\Delta$.

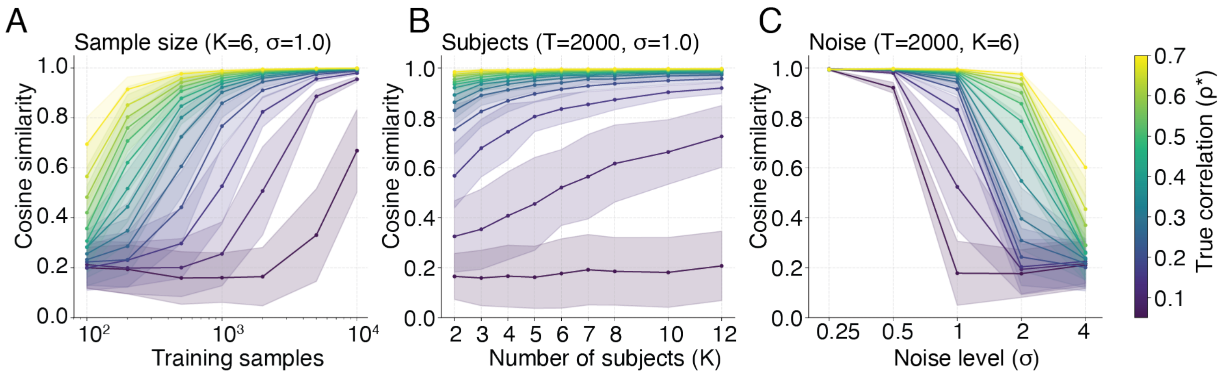


Figure S3: **Sensitivity of component recovery to sample size, number of subjects, and noise level.** Each panel shows cosine similarity between true and recovered behavioral projections for 14 components with target correlations from 0.05 to 0.70 (color scale), with mean ± std across 100 repetitions. The noise level $\sigma$ is the per-feature standard deviation of the i.i.d. Gaussian noise in the synthetic generative model (Supplementary Note 3). **(A)** Recovery as a function of training sample size ($K = 6$, $\sigma = 1.0$). Components with $\rho \geq 0.5$ are reliably recovered from 500 samples. Very weak components ($\rho \leq 0.05$) remain unreliable even with 10,000 samples. **(B)** Recovery as a function of the number of subjects ($T = 2{,}000$, $\sigma = 1.0$). Performance improves with additional subjects, with diminishing returns beyond six for moderate-to-strong components. **(C)** Recovery as a function of noise level ($T = 2{,}000$, $K = 6$). Strong components tolerate substantial noise ($\sigma \leq 2.0$). Weak components degrade rapidly above $\sigma = 1.0$.

Figure S4: **Shared behavioral weight profiles for the center-out (CO) task.**

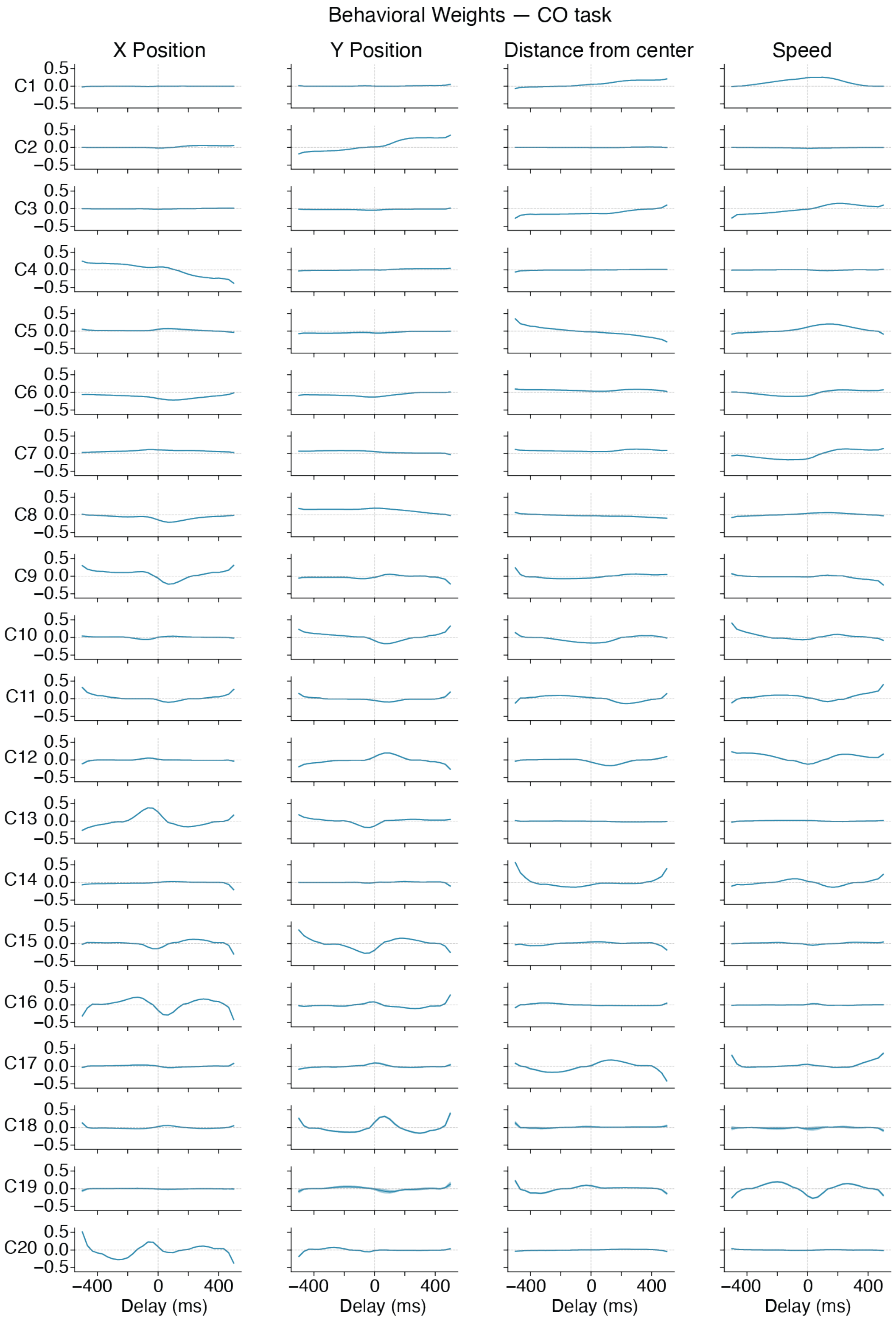

Figure S4: **(continued)** Each row shows the fold-averaged shared behavioral projection $\mathbf{b}$ for one of 20 ShaReD components (C1–C20), with columns corresponding to four kinematic variables, namely $x$ position, $y$ position, distance from workspace center, and speed. Weights are plotted as a function of temporal delay ($-500$ to $+500$ ms relative to neural activity). Component 1 is a radial signal dominated by distance from center and speed, with $x$ and $y$ position weights near zero, consistent with its direction-invariant tuning in Fig. 2E. Component 2 is dominated by $y$ (vertical) position. Subsequent components capture progressively more complex mixtures of position, speed, and distance across temporal offsets. Weights are distributed across a range of lags rather than a single delay, and the grouped smoothness penalty renders these temporal profiles continuous. Shaded bands indicate SEM across 10 cross-validation folds. $n = 73$ sessions from 3 monkeys.

Figure S5: **Shared behavioral weight profiles for the random target (RT) task.**

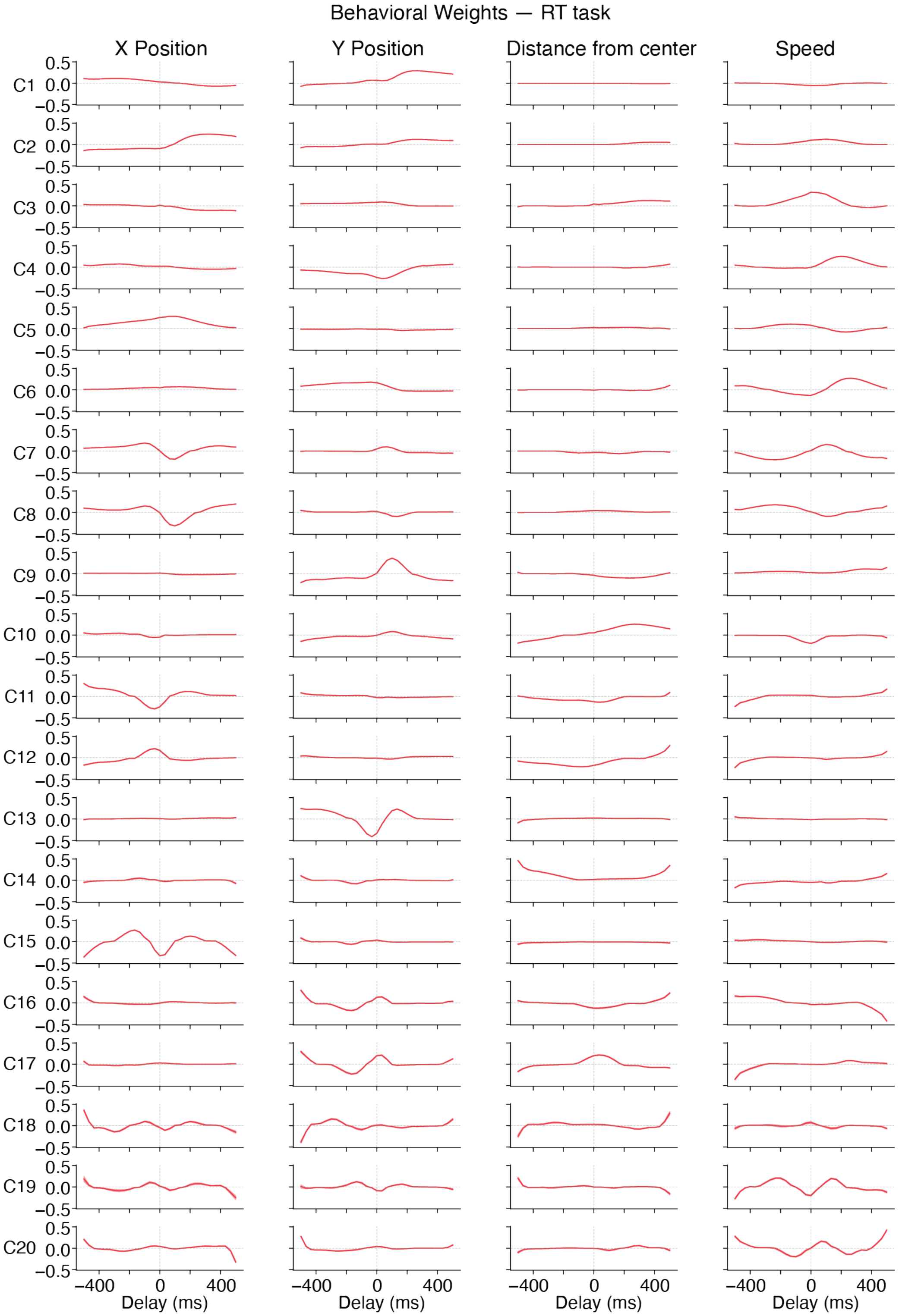

Figure S5: **(continued)** Each row shows the fold-averaged shared behavioral projection $\mathbf{b}$ for one of 20 ShaReD components (C1–C20), with columns corresponding to four kinematic variables, namely $x$ position, $y$ position, distance from workspace center, and speed. Weights are plotted as a function of temporal delay ($-500$ to $+500$ ms relative to neural activity). The first several components show interpretable structure, with Component 1 primarily depending on position, and subsequent components also reflecting speed and distance at varying lags. Both projections are normalized, so these panels show how each component distributes a fixed norm across features and lags rather than how the components differ in overall magnitude. RT components reach lower per-component correlations than CO (Fig. 2B), consistent with the task's higher behavioral variability. Despite different movement statistics (random targets versus eight fixed directions), similar temporal patterns emerge across both tasks. The best cross-task cosine similarity matches exceed 0.70 (Fig. S6). Shaded bands indicate SEM across 10 cross-validation folds. $n = 26$ sessions from 3 monkeys.

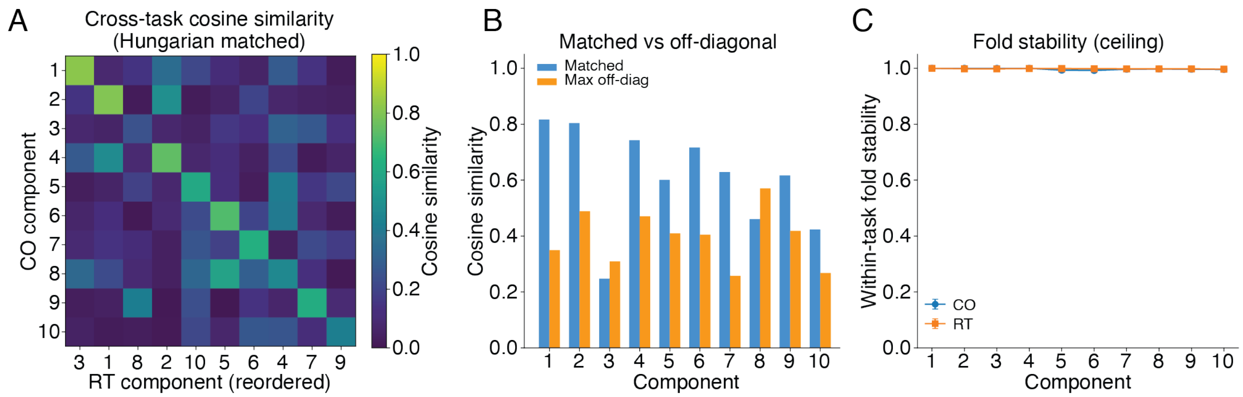


Figure S6: **Cross-task behavioral weight similarity. (A)** Cosine similarity matrix between CO and RT behavioral projections $\mathbf{b}$ for the first 10 components, with RT components reordered by Hungarian matching so that matched pairs lie on the diagonal. Because the deflation-based extraction order differs between tasks, each matched RT component carries a different index from its CO partner ($x$-axis labels). For example, CO Component 1 matches RT Component 3 at cosine similarity 0.82, CO Component 2 matches RT Component 1 at 0.80, and CO Component 4 matches RT Component 2 at 0.74. **(B)** Matched (diagonal) versus maximum off-diagonal cosine similarity per component. Matched similarity exceeds 0.7 for components 1, 2, 4, and 6, and is lowest for component 3 ($\sim$0.25), consistent with the component-specific transfer pattern in Fig. 3B. **(C)** Within-task across-fold stability of $\mathbf{b}$ vectors over 10 cross-validation folds. The shared projections are highly reproducible (all $>$0.99 for the first 10 components in both tasks). This stability ceiling is far above the cross-task similarity, indicating that cross-task differences reflect genuine task effects rather than estimation noise. Error bars indicate SD across the 45 fold pairs.

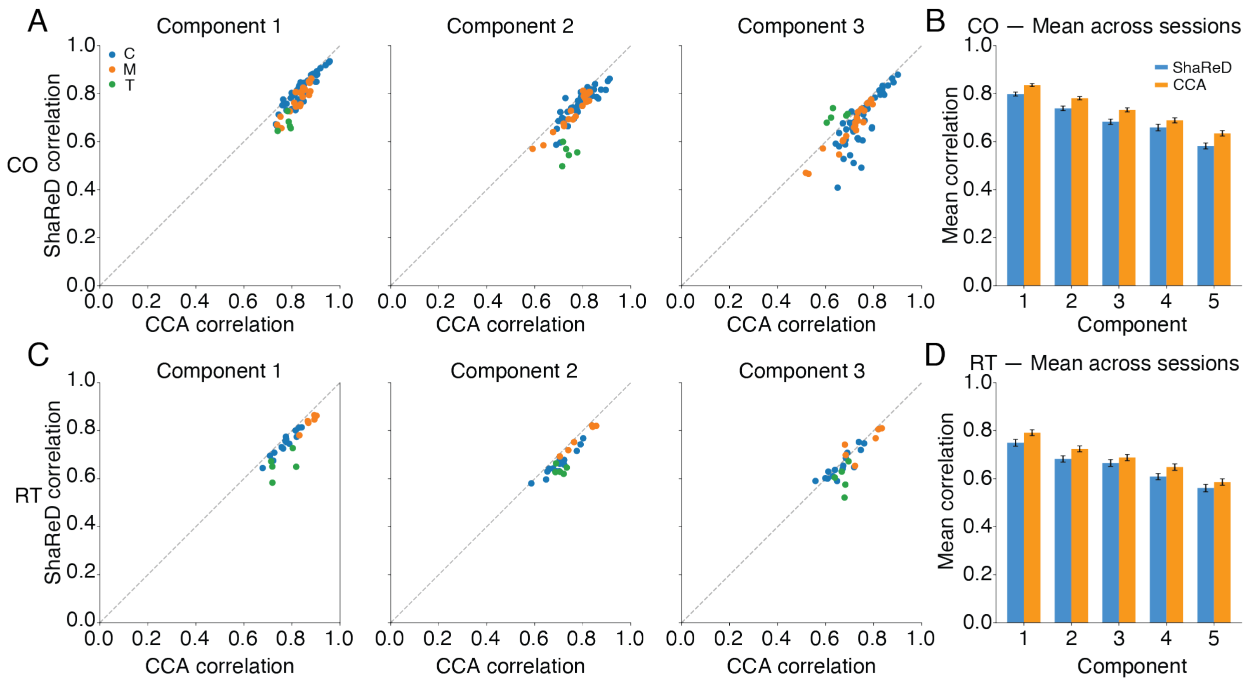


Figure S7: **Within-session CCA baseline comparison.** Per-session cross-validated correlations for ShaReD versus unconstrained within-session CCA for the CO task (A, B) and RT task (C, D). Each dot represents one session, colored by monkey (C in blue, M in orange, T in green). **(A)** Components 1–3 shown individually. Most sessions fall near or slightly below the identity line, with CCA achieving modestly higher per-session correlations (∼5% for Component 1). The gap reflects the cost of ShaReD's common-**b** constraint, since CCA fits an independent behavioral projection per session and gains per-session accuracy at the expense of cross-session comparability. **(B)** Mean correlation across the first 5 components for ShaReD and CCA. The modest CCA advantage indicates that the shared constraint does not substantially sacrifice per-session encoding quality, while providing a common behavioral reference frame across all sessions. **(C, D)** As (A, B) but for the RT task. The same modest CCA advantage holds, with comparable gap sizes across components. Error bars in (B) and (D) indicate SEM across sessions. $n = 73$ CO sessions, $n = 26$ RT sessions.

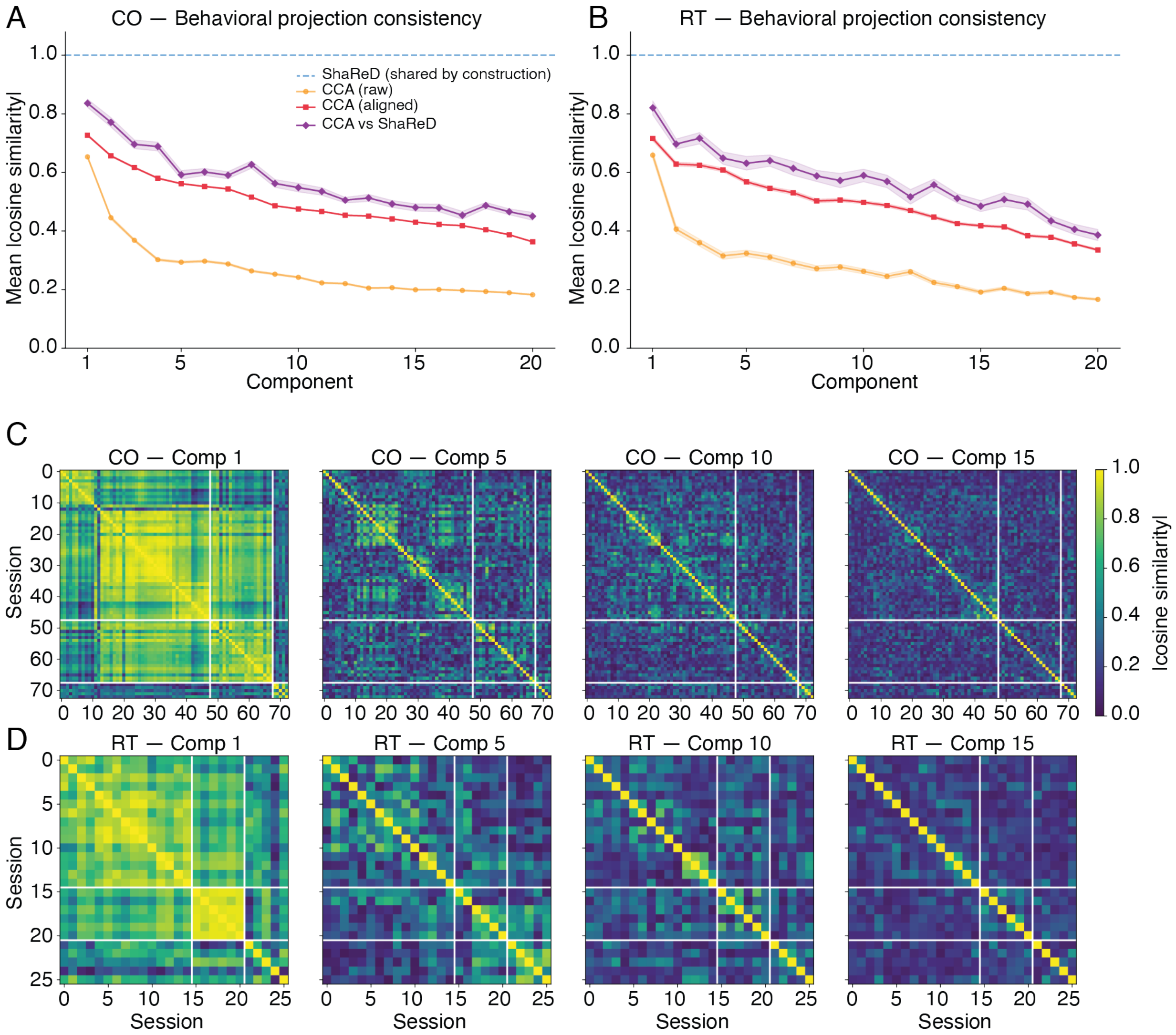


Figure S8: **CCA behavioral projection consistency across sessions.** **(A, B)** Mean absolute cosine similarity of behavioral projections across sessions as a function of component number. ShaReD projections (blue dashed, = 1.0 by construction) are compared with per-session CCA projections under three conditions, namely raw cosine similarity of CCA components (orange, no alignment), aligned via Hungarian matching with optimal sign correction (red), and CCA-versus-ShaReD similarity (purple). Even after optimal post-hoc alignment, per-session CCA projections show limited consistency (CO ∼0.73 for Component 1 and RT ∼0.72, with means across 20 components of 0.50 and 0.49 respectively), indicating that independent per-session fitting does not recover ShaReD's common reference frame. The CCA-versus-ShaReD similarity (purple) exceeds the CCA self-consistency (red), suggesting that ShaReD captures a meaningful average of the per-session solutions. **(C, D)** Pairwise cosine similarity heatmaps for selected CCA components (1, 5, 10, 15), with sessions grouped by monkey (white lines). Component 1 shows moderate within-monkey consistency but lower across-monkey agreement. Later components show near-diagonal-only structure, indicating that per-session CCA projections are largely session-specific. Shaded bands in (A) and (B) indicate SEM across sessions.

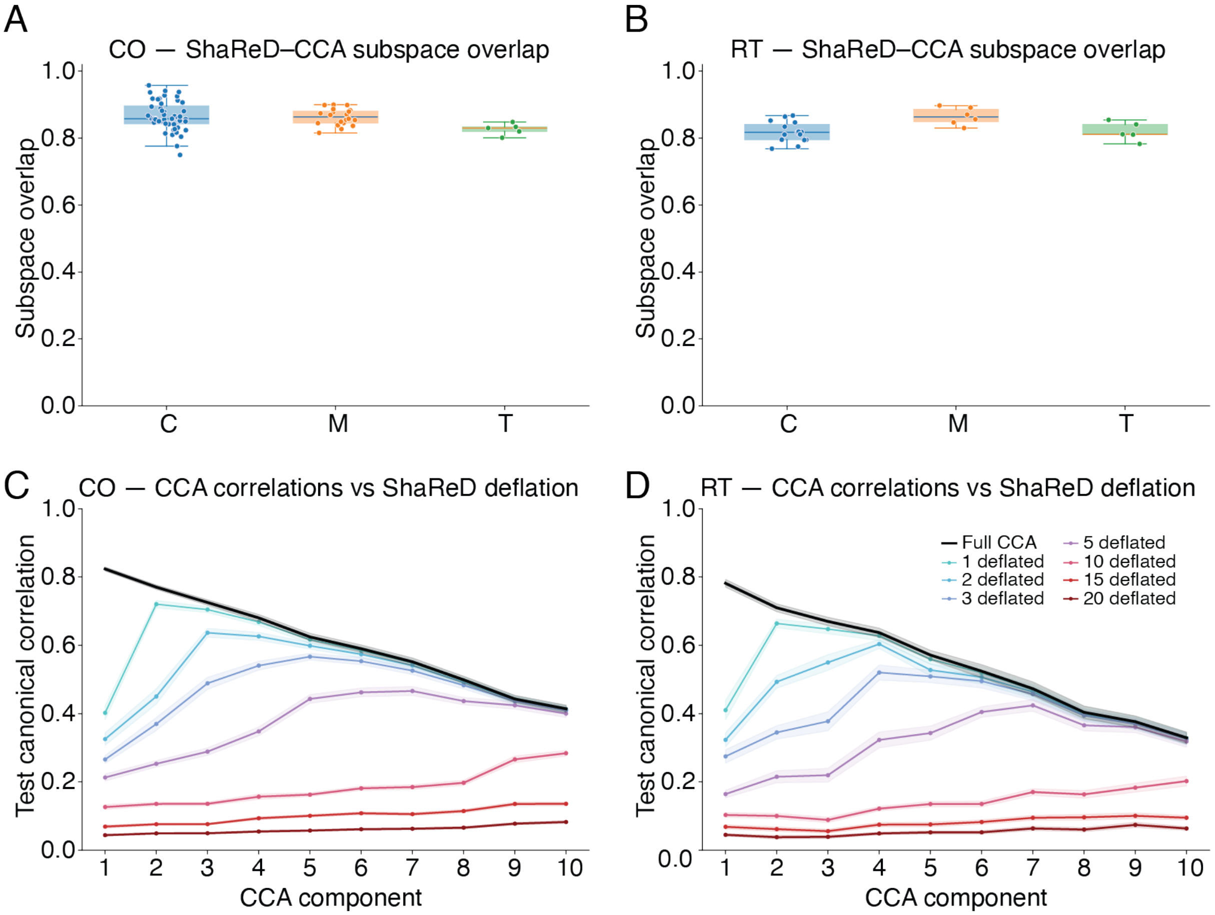


Figure S9: **CCA-ShaReD subspace overlap and deflation test. (A, B)** Subspace overlap between the ShaReD-identified subspace and the per-session CCA subspace for each monkey, measured as the mean absolute cosine similarity between the ShaReD **b** vector and the span of the top-20 CCA canonical variates. Overlap is high (CO 0.74–0.94 across sessions, mean 0.86 and RT 0.77–0.90, mean 0.83), indicating that ShaReD and CCA identify similar behavioral subspaces despite their different optimization criteria. Boxes show the interquartile range with the median, and whiskers extend to 1.5 times the IQR. **(C, D)** CCA test canonical correlations before and after ShaReD deflation. Black line shows the original (undeflated) CCA correlations across 10 canonical components. Colored lines show correlations after deflating 1–20 ShaReD components. Deflating even a few ShaReD components substantially reduces CCA correlations, showing that ShaReD captures the dominant neural-behavioral variance that CCA also identifies. This is stronger evidence than geometric overlap alone, since two subspaces can be aligned without sharing the covariance that produces the correlation. Deflating the ShaReD components removes the covariance CCA's correlations are built on, so the correlations drop. Shaded bands in (C) and (D) indicate SEM across sessions.

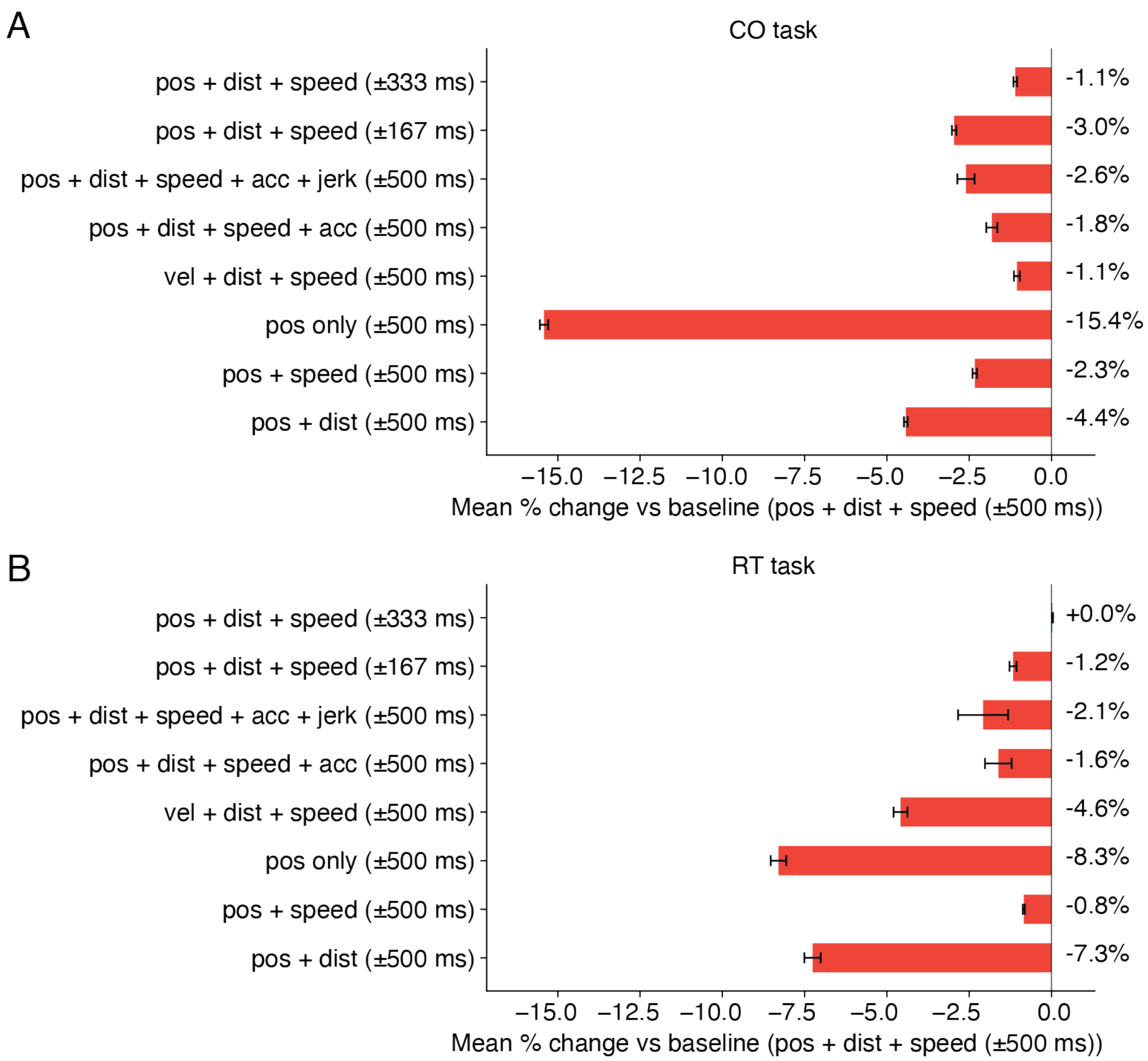


Figure S10: **Feature set ablation.** Mean percent change in cross-validated test-set correlation relative to the selected baseline feature set (position + distance + speed ±500 ms temporal window) for the CO task (A) and RT task (B). The full feature-selection sweep tested 63 candidate configurations. This figure shows representative ablations around the selected feature set. Tested alternatives include reduced temporal windows (±333 ms, ±167 ms), additional kinematic derivatives (acceleration, jerk), removal of individual features, reduction to position only, and substitution of velocity for position. All three kinematic variables contributed, with the dominant variable differing by task. Using position alone reduces performance by 15.4% in CO and 8.3% in RT relative to the selected feature set, indicating that distance and speed together contribute substantially. Distance contributes more in CO (−2.3%) than RT (−0.8%), while speed contributes more in RT (−7.3%) than CO (−4.4%). Replacing position with velocity reduces performance by 1.0% in CO and 4.6% in RT. Adding acceleration or jerk provides no additional benefit and slightly reduces performance (acceleration $\leq 1.8\%$, acceleration plus jerk $\leq 2.6\%$), consistent with the extra features raising behavioral dimensionality without contributing shared structure. Reducing the temporal window from ±500 ms to ±333 ms reduces performance by up to $\sim$1% (CO 1.1%, RT $\sim$0%), while reducing it to ±167 ms reduces performance by approximately 3.0% in CO and 1.2% in RT. Error bars show SEM across cross-validation folds. $n = 73$ CO sessions, $n = 26$ RT sessions.

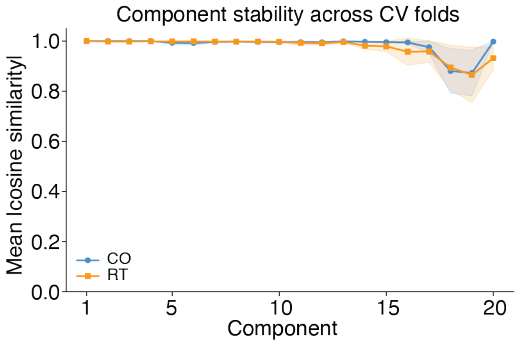


Figure S11: **Stability of shared behavioral projections across cross-validation folds.** Mean absolute cosine similarity of the shared behavioral projection **b** between all pairs of 10 cross-validation folds, shown for all 20 components. CO task (blue) and RT task (orange) both show near-perfect stability ($> 0.99$) for the first $\sim 12$ components, with modest degradation for later components (minimum $\sim 0.87$). Shaded regions indicate $\pm 1\,\mathrm{SD}$ across the 45 fold pairs. The high stability indicates that ShaReD's common projections are reproducible and not sensitive to the particular data partition, supporting the use of fold-averaged weights for interpretation (Figs. S4–S5).

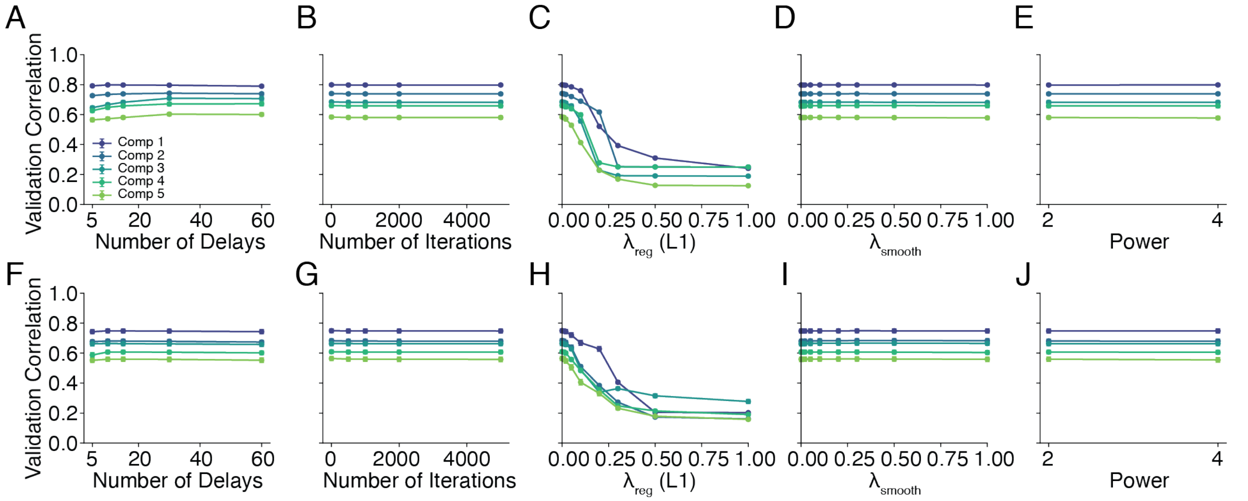


Figure S12: **Hyperparameter sensitivity.** One-at-a-time sweep of five ShaReD hyperparameters, showing the per-component cross-validated validation correlation (colored lines, 5 components) for the CO task (A–E) and RT task (F–J). Hyperparameters were selected on a nested split into sub-training, validation, and test partitions, and the curves plot the validation correlation that drives this selection. All other parameters are held at their defaults (number of delays = 15, iterations = 1000, $\lambda_{\text{reg}} = 0.01$, $\lambda_{\text{smooth}} = 0.01$, power = 2). **(A)** Number of temporal delays (5–60, spanning $\pm 167$ to $\pm 2000$ ms). Performance is stable across the tested range, with mild improvement at 30 delays. **(B)** Number of optimization iterations (0–5000). The closed-form initialization (0 iterations) already achieves near-optimal performance. Iterative refinement provides modest gains that plateau by 1000 iterations. **(C)** L1 regularization strength $\lambda_{\text{reg}}$ (0–1). Performance is stable for small penalties and declines as $\lambda_{\text{reg}}$ increases, since the proximal soft-threshold followed by renormalization to unit norm progressively drives $\mathbf{b}$ toward a single feature, removing the multi-lag structure the leading components rely on. **(D)** Smoothness regularization strength $\lambda_{\text{smooth}}$ (0–1). Performance is stable across the range, with slight improvement at moderate values. **(E)** Objective function power (2 versus 4). The squared objective ($p = 2$) and quartic objective ($p = 4$) produce comparable results for the first four components. The fifth component shows mild degradation under the quartic objective. **(F–J)** As (A–E) but for the RT task. Sensitivity patterns match the CO results, except in (J), where both the fourth and fifth components show mild degradation under the quartic objective. Error bars indicate SEM across sessions. $n = 73$ CO sessions, $n = 26$ RT sessions.

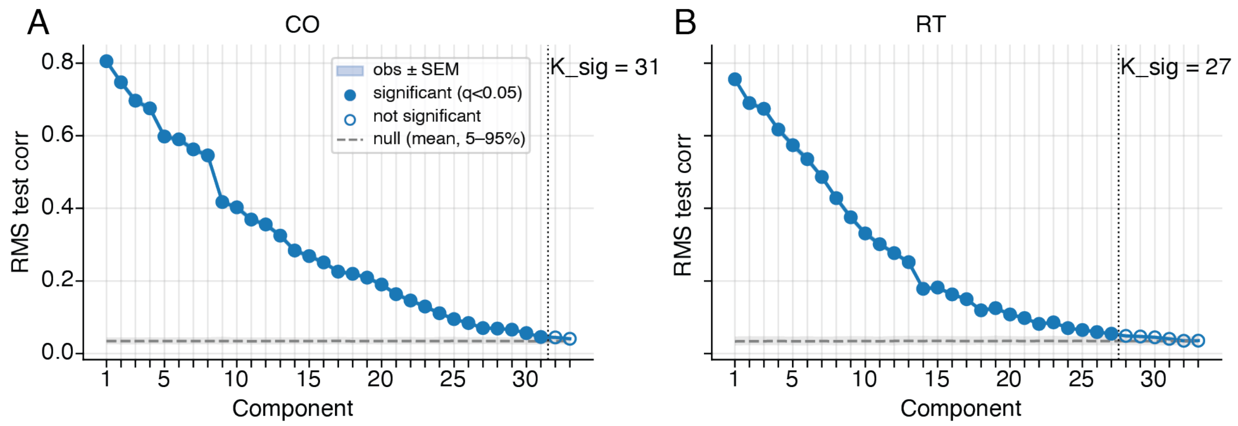


Figure S13: **Per-component significance for the monkey motor cortex analysis.** Observed RMS cross-validated neural-behavioral correlation for each ShaReD component (points, with ± SEM shaded) against the refit-shuffle null distribution (gray dashed line for the null mean, shaded band for 5–95% of the null) for the center-out (A) and random target (B) tasks. The null is obtained by refitting ShaReD on shuffled behavior. A component is significant when its observed correlation exceeds the null at a Benjamini–Hochberg false discovery rate (BH-FDR) of $q < 0.05$. Filled markers denote significant components and open markers denote non-significant ones. The vertical dotted line marks the end of the leading contiguous run of significant components, whose length is reported as $K_{\text{sig}}$ ($K_{\text{sig}} = 31$ for CO, $K_{\text{sig}} = 27$ for RT). The main-text analyses interpret only the leading 20 components, a conservative cap below the full significant run. $n = 73$ CO sessions, $n = 26$ RT sessions from 3 monkeys.

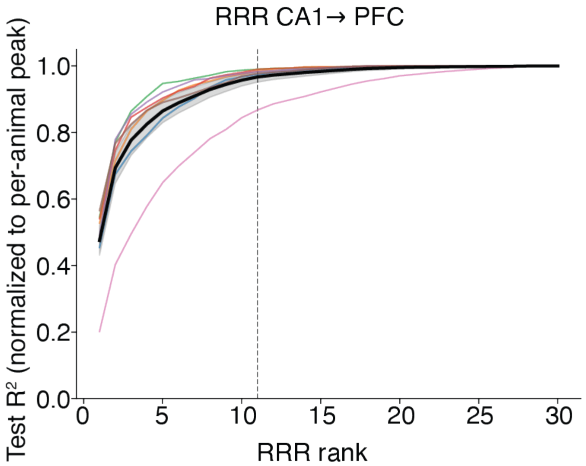


Figure S14: **Reduced-rank regression rank selection for the rat CA1→PFC mapping.** Per-animal 5-fold cross-validated CA1→PFC test $R^2$ as a function of reduced-rank regression rank, with one curve per rat. The black curve shows the mean across rats, and the shaded band around it indicates SEM across rats. For each animal, rank was selected by the one-standard-error (1-SE) rule, which takes the smallest rank whose cross-validated $R^2$ is within one standard error across folds of the best-performing rank (full procedure in Methods). This gives per-animal rank estimates with a median of 11 and a range of 5–15 across the 7 rats. The fixed rank of 11 (dotted line) used for all animals throughout the main analysis falls at this median. $n = 7$ rats.

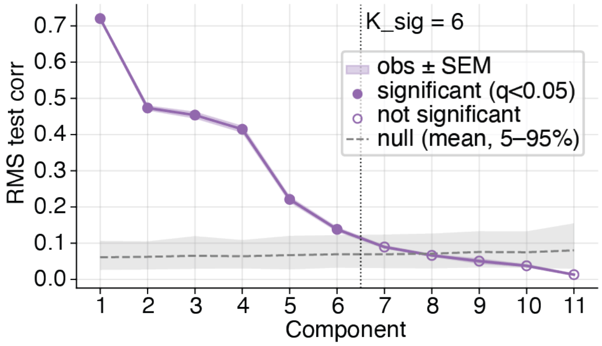


Figure S15: **Per-component significance for the rat CA1→PFC analysis.** Observed RMS cross-validated neural-behavioral correlation for each of the 11 ShaReD components against the refit-shuffle null distribution (gray). Significance was assessed against this null with Benjamini–Hochberg FDR correction at $q < 0.05$ (see Methods). Six of the 11 components are significant, with $p_{\mathrm{FDR}} = 0.004$ for components 1–5 and $p_{\mathrm{FDR}} = 0.040$ for component 6. Components 7–11 do not reach significance. $n = 7$ rats.

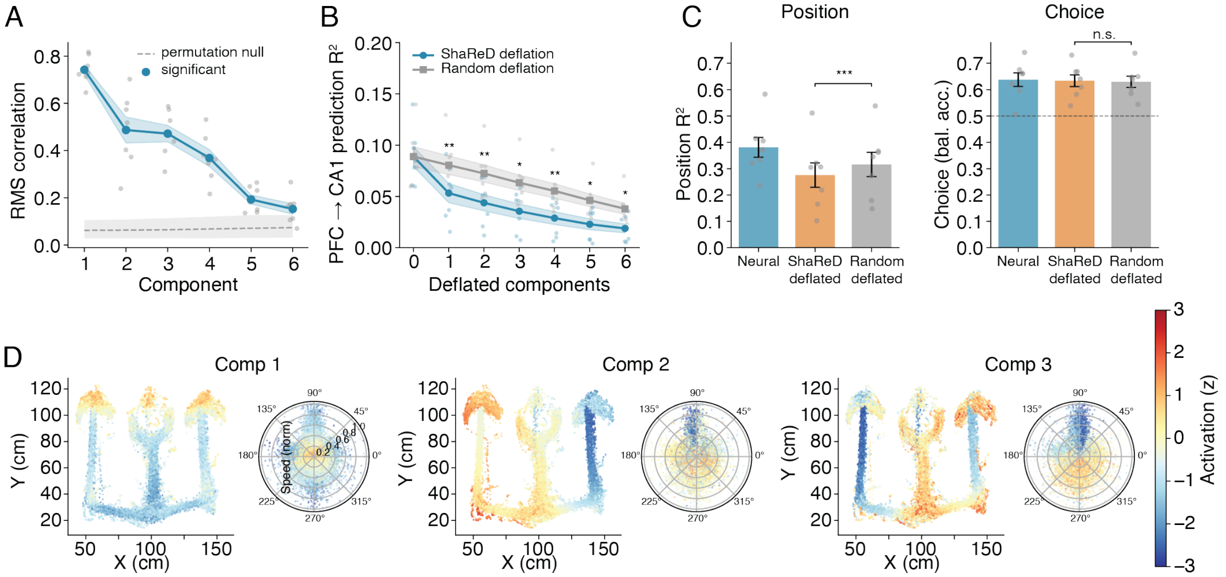


Figure S16: **Reversed-direction communication subspace analysis (PFC→CA1).** Same format as Fig. 4 but with the prediction direction reversed. PFC activity is used to predict CA1 activity via reduced-rank regression, and ShaReD identifies shared behavioral dimensions within the PFC→CA1 communication subspace. **(A)** RMS correlation between neural and behavioral projections for 6 ShaReD components extracted from the PFC→CA1 communication subspace. Correlations decline with component number, following a similar pattern to the CA1→PFC direction. **(B)** PFC→CA1 prediction $R^2$ as a function of the number of deflated ShaReD components (blue) versus random deflation of matched dimensionality (gray). ShaReD deflation reduces prediction more rapidly than random deflation, indicating that the identified components carry specifically behavioral variance. Shaded bands indicate SEM across 7 rats. **(C)** Position decoding ($R^2$, left) and choice classification (balanced accuracy, right) from the full PFC population (blue), after ShaReD deflation (orange), and after random deflation (gray). Choice labels are the upcoming left versus right turn, decoded with balanced class weights and scored by balanced accuracy, so chance is 0.5 regardless of any imbalance in label counts (Methods). ShaReD deflation significantly reduces position decoding. Choice decoding shows a small numerical reduction after ShaReD deflation that does not reach significance after Bonferroni correction, as in the CA1→PFC direction (compare Fig. 4E), so the dissociation between position and choice holds in both directions. Error bars indicate SEM across 7 rats. **(D)** Spatial and directional tuning of the first three ShaReD components, shown as position maps (colored by component activation) and polar head-direction tuning plots for an example animal (ER1). $n = 7$ rats.
